# A Multimodal Characterization of Low-Dimensional Thalamocortical Structural Connectivity Patterns

**DOI:** 10.1101/2024.02.01.578366

**Authors:** Alexandra John, Meike D. Hettwer, H. Lina Schaare, Amin Saberi, Şeyma Bayrak, Bin Wan, Jessica Royer, Boris C. Bernhardt, Sofie L. Valk

## Abstract

The human thalamus is a bilateral and heterogeneous grey matter structure that plays a crucial role in coordinating whole-brain activity. Investigations of its complex structural and functional internal organization revealed to a certain degree overlapping parcellations, however, a consensus on thalamic subnuclei boundaries remains absent. Recent work suggests that thalamic organization might additionally reflect continuous axes transcending nuclear boundaries. In this study, we used a multimodal approach to uncover how low-dimensional axes that describe thalamic connectivity patterns to the cortex are related to internal thalamic microstructural features, functional connectivity, and structural covariance. We computed a thalamocortical structural connectome via probabilistic tractography on diffusion MRI and derived two main axes of thalamic organization. The principal thalamic gradient, extending from medial to lateral and differentiating between transmodal and unimodal nuclei, was related to intrathalamic myelin profiles, and patterns of functional connectivity, while the secondary axis showed correspondence to core-matrix cell type distributions. Lastly, exploring multimodal thalamocortical associations on a global scale, we observed that the medial-to-lateral gradient consistently differentiated limbic, frontoparietal, and default mode network nodes from dorsal and ventral attention networks across modalities. However, the link with sensory modalities varied. In sum, we show the coherence between lower dimensional patterns of thalamocortical structural connectivity and various modalities, shedding light on multiscale thalamic organization.

## 1. Introduction

As part of the diencephalon, the human thalamus is a bilateral grey matter structure that orchestrates whole-brain activity (Shine et al., 2023). Due to its extensive connections to the entire cerebral cortex, and subcortical structures such as the basal ganglia and the cerebellum, the thalamus can be characterized as a ‘connector hub’ region in the brain. In the past, the thalamus was primarily described as a relay passively transferring information to the cortex. More recent perspectives, however, are superseding this cortico-centric notion and emphasize the thalamic impact on mediating the corticocortical information transfer, the integration of information between cortical networks (Sherman, 2016; Hwang et al., 2017; Sherman and Guillery, 2013) and its role in cognition (Hwang et al., 2022; Halassa and Kastner, 2017; Schiff, 2008; de Bourbon-Teles et al., 2014; Saalmann et al., 2012).

The internal organization of the thalamus is highly complex. *Ex vivo*, it has been parcellated into multiple subnuclei through analyzing histologically stained dissected *post-mortem* brains (Jones, 1985; Morel et al., 1997). These thalamic subnuclei contain a blend of two thalamic cell types, referred to as ‘core’ and ‘matrix’ cells, in varying proportions (Clascá et al., 2012; Jones, 1998). While core cells tend to target specifically layer IV and V of primary sensory cortex, matrix cells have widespread cortical projections and innervate the supragranular layers I-III (Clascá et al., 2012; Jones, 2009, 2001, 1998). Further, the thalamic nuclei can be broadly classified into first-order nuclei (*i.e.*, ventral posterolateral nucleus) that receive ascending sensory and modulatory cortical input, and higher-order nuclei (*i.e.*, mediodorsal nucleus) that receive their input entirely from the cortex (Sherman, 2012; Sherman and Guillery, 1998). Compared to *post-mortem* studies, non-invasive neuroimaging using magnetic resonance imaging (MRI) provides the opportunity to study the thalamic organization and its relationship to the cortex *in vivo*, enabling data acquisition in large sample sizes and investigations of structure-function coupling. While delineating the thalamus locally using standard T1- and T2-weighted MRI images remains challenging due to poor tissue contrast (Tourdias et al., 2014), utilizing global approaches, such as structural and functional connectivity methods, that consider the extensive interrelation between the thalamus and the cerebral cortex has provided valuable insights into thalamic organization (Behrens et al., 2003a; Ji et al., 2016). In the pioneering work of Behrens et al. (2003a), the thalamus was parcellated based on thalamocortical (TC) probabilistic tractography using diffusion weighted imaging (DWI). This parcellation was highly inter- and intra-subject reproducible (Traynor et al., 2010) and showed robust correspondence with thalamic function (Johansen-Berg et al., 2005). Further approaches based on DWI were explored to uncover thalamic organization (O’Muircheartaigh et al., 2011; Battistella et al., 2017; Stough et al., 2014) and accordingly several thalamic atlases have been published (Behrens et al., 2003a; Iglesias et al., 2018; Najdenovska et al., 2018). Alongside structural connectivity, the functional coupling between the thalamus and cortex has been used to gain insights into the thalamic organization through TC functional connectivity (Ji et al., 2016; Hwang et al., 2017; Zhang et al., 2008). Taken together, the existing literature suggests that the heterogeneous organization of the thalamus can be subdivided into nuclei dependent on scale and modality. These subnuclei vary in morphology, connectivity, and function.

Though subregions derived from different modalities overlap to a certain degree, a consensus of thalamic parcellation remains absent due to the high complexity of this region (Iglehart et al., 2020). Specifically, drawing clear and universal boundaries proves challenging because of the complex connectivity profiles of thalamic nuclei that target multiple cortical regions and are associated with various functional networks (Hwang et al., 2017). From a developmental point of view, the emergence of thalamic structural and functional properties is guided by molecular gradients of morphogens and transcription factors (López-Bendito and Molnár, 2003; Vogel et al., 2022; Govek et al., 2022). This is reflected in transitional patterns of gene expression and cytoarchitecture transcending hard borders of thalamic nuclei (Roy et al., 2022; Gao et al., 2020; Phillips et al., 2019). Indeed, applying transcriptional profiling in mice revealed that gradual changes in gene expression are tied to anatomical and electrophysiological properties (Phillips et al., 2019). In line with this, recent advances of studying brain organization have shifted their focus on revealing spatially graded changes of neurobiological properties across the brain, in addition to the traditional approaches of defining discrete brain regions (Bernhardt et al., 2022; Margulies et al., 2016; Smallwood et al., 2021; Valk et al., 2020; Paquola et al., 2019; Huntenburg et al., 2018). These continuous axes of spatial variation are referred to as ‘gradients’. While this approach has mainly been applied to understand macroscale cortical organization, recent work uncovered transitional axes that help explain organizational patterns of the human thalamus (Oldham and Ball, 2023; Zheng et al., 2023; Yang et al., 2020). Based on the joint analysis of TC structural connectivity and gene expression data, a phylogenetically conserved medial-to-lateral axis has been reported that captured transitions in cell type variations (Oldham and Ball, 2023). Furthermore, TC functional connectivity has been shown to follow a medial-to-lateral axis associated with thalamic grey matter volume, and an anterior-to-posterior axis corresponding to functional networks (Yang et al., 2020). These axes might arise from smooth transition at the microscale level (Phillips et al., 2019; Roy et al., 2022).

On top of genetic determination, thalamic functional activity has a substantial influence on cortical maturation through activity-dependent maturational processes (Antón-Bolaños et al., 2018; López-Bendito, 2018). A global measure used in neuroimaging that is hypothesized to capture both genetic and maturational coherences is structural covariance, which describes covariation in structural properties, such as cortical thickness, across different brain regions (Alexander-Bloch et al., 2013a). Although the biological mechanisms driving structural covariance remain incompletely understood, they may implicate activity-induced synaptogenesis and/or synchronous neurodevelopment (Alexander-Bloch et al., 2013b; Evans, 2013), and relate to shared genetic effects (Valk et al., 2020; Schmitt et al., 2009). In mice, TC structural covariance has been used to investigate thalamic organization, which showed some correspondence between TC structural and functional connectivity (Yee et al., 2024), suggesting a link between shared maturational patterning and connectivity in the thalamus.

In the current study, we explored how the internal organization of the human thalamus based on its structural connections to the cortex corresponds with the distribution of thalamic microstructural features, as well as TC functional connectivity and structural covariance. We assessed the structural connections between thalamic-seeds and the cortex by computing probabilistic tractography and extracted low-dimensional axes of thalamic organization. To contextualize our findings, we spatially associated these axes to intrathalamic microstructural features, such as grey matter myelin based on quantitative T1 (qT1), and the distribution of cell types based on gene expression data. Additionally, we explored the association between the lower dimensional organization of structural and functional connectivity. Moving to a more global perspective, we further investigated how the thalamic gradients are related to macroscale cortical patterns and studied the link to TC functional connectivity and qT1- based TC structural covariance.

## 2. Results

### Thalamic Gradients Based on TC Structural Connectivity (Figure 1)

The first aim of this study was to investigate the spatial organization of the thalamus based on TC structural connectivity. To do so, we applied probabilistic tractography to map the white matter connections between thalamic voxels and 100 ipsilateral cortical parcels in each subject. The resultant thalamic-seed-by-cortical-parcel structural connectivity matrices contained the number of streamlines between seeds and parcels. After averaging across subjects, the group-level structural connectivity matrix was normalized column-wise (Figure 1A). To uncover the lower dimensions of this structural connectivity matrix, we first derived an affinity matrix (Figure 1B) and further applied dimensionality reduction via diffusion map embedding. The decomposition yielded ten unitless gradient components, where nodes that share similar connectivity profiles were embedded closer together, and nodes with little connectivity similarity mapped further apart. Each gradient component represented a particular axis of the thalamic organization and explained a certain amount of variance (Figure 1C). Henceforth, we focused on the first two components/gradients that accounted for a total variance of 46.42 % in the left hemisphere and 46,47 % in the right hemisphere. The principal TC structural connectivity gradient (G1_sc_; explained variance left hemisphere (LH): 25.99 %, right hemisphere (RH): 25.82 %) defined a medial to lateral-central axis of the thalamus (Figure 1D). The secondary gradient (G2_sc_; explained variance LH: 20.43 %, RH: 20.65 %) located one apex at the medial-anterior but also posterior pole of the thalamus and the opposite apex intersected the thalamus from anterior-lateral to medial-central (Figure 1E). Further, we showed robustness of gradient patterns derived from structural connectivity matrices thresholded at different percentiles (Supplementary Figure 1).

**Figure 1:**
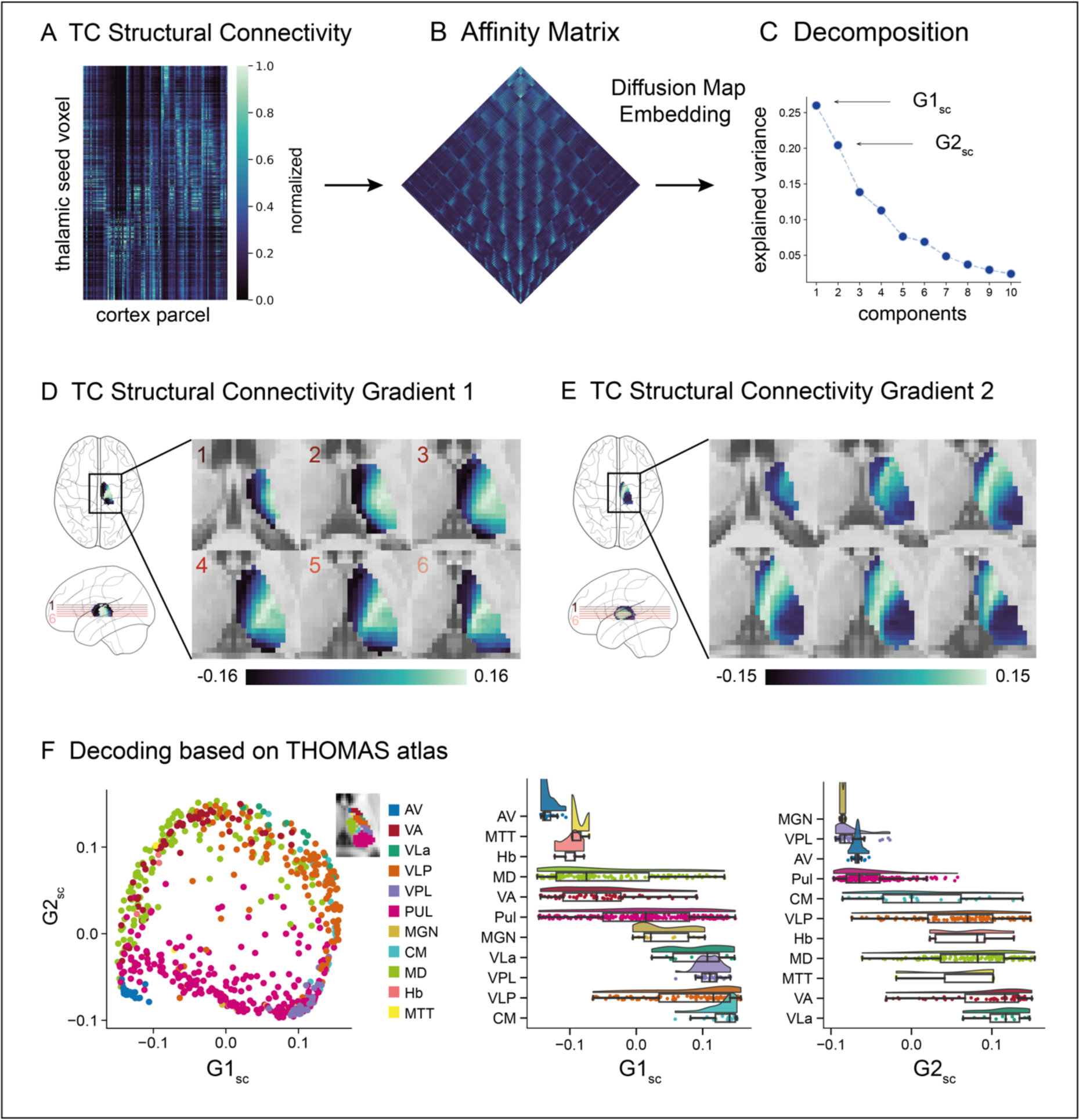
Thalamocortical Structural Connectivity Gradients. A. Normalized group-level structural connectivity matrix resulting from probabilistic tractography computation between thalamic seed-voxels and cortical parcels (*i.e*., TC). **B** Affinity matrix derived from group-level structural connectivity matrix using a normalized angle similarity kernel. **C** Decomposition of affinity matrix into ten gradient components using diffusion map embedding. For each component the corresponding explained variance is displayed. **D** Gradient loadings of component 1 (G1_sc_) projected on the thalamus (axial planes). The red lines in the glass brain indicate the position of each respective axial slice of the displayed thalamus. **E** Gradient loadings of component 2 (G2_sc_) projected on the thalamus. Slice positions are congruent to D. **F** Left: Decoding of G1_sc_ and G2_sc_ based on THOMAS atlas. 2D space framed by G1_sc_ and G2_sc_, color-coded by thalamic subnuclei. Middle and right: Raincloud plots display the gradient loadings of G1_sc_ and G2_sc_ per nucleus and are ordered by median, respectively. All results are presented for the left hemisphere, however, they were similarly replicated in the right thalamus (Supplementary Figure 2). Abbreviations in **F**: AV: Anterior ventral nucleus, VA: Ventral anterior nucleus, VLa: Ventral lateral anterior nucleus, VLP: Ventral lateral posterior nucleus, VPL: Ventral posterior lateral nucleus, Pul: Pulvinar nucleus, MGN: Medial geniculate nucleus, CM: Centromedian nucleus, MD: Mediodorsal nucleus, Hb: Habenular nucleus, MTT: Mammillothalamic tract

To relate the TC structural connectivity gradients to previously defined discrete thalamic subnuclei, we used the THOMAS parcellation in MNI152 space (Su et al., 2019; Saranathan et al., 2021). In the 2D gradient frame, we color-coded datapoints according to the corresponding subnucleus showing the spatial distribution of subnuclei along these two axes. Ordering the distinct nuclei based on the median of their gradient loadings revealed for G1_sc_ the following order: AV, MTT, Hb, MD, VA, Pul, MGN, VLa, VPL, VLP, CM^1^ and hence differentiated between higher-order nuclei and sensorimotor nuclei. Ordering the nuclei along G2_sc_ resulted in: MGN, VPL, AV, Pul, CM, VLP, Hb, MD, MTT, VA, VLa^1^, and henceforth does not separate first-order and higher-order nuclei (Figure 1F). Here, results of the left hemisphere are reported, however, they were similarly replicated in the right hemisphere (Supplementary Figure 2).

### TC Structural Connectivity Gradients are Associated with Microstructure and Functional Connectivity (Figure 2)

After computing structural connectivity gradients, we probed whether they relate to underlying thalamic organizational properties, such as microstructural features (distribution of qT1 values as a proxy for myelin, distribution of core and matrix cells), and thalamic gradients based on TC functional connectivity.

**Figure 2:**
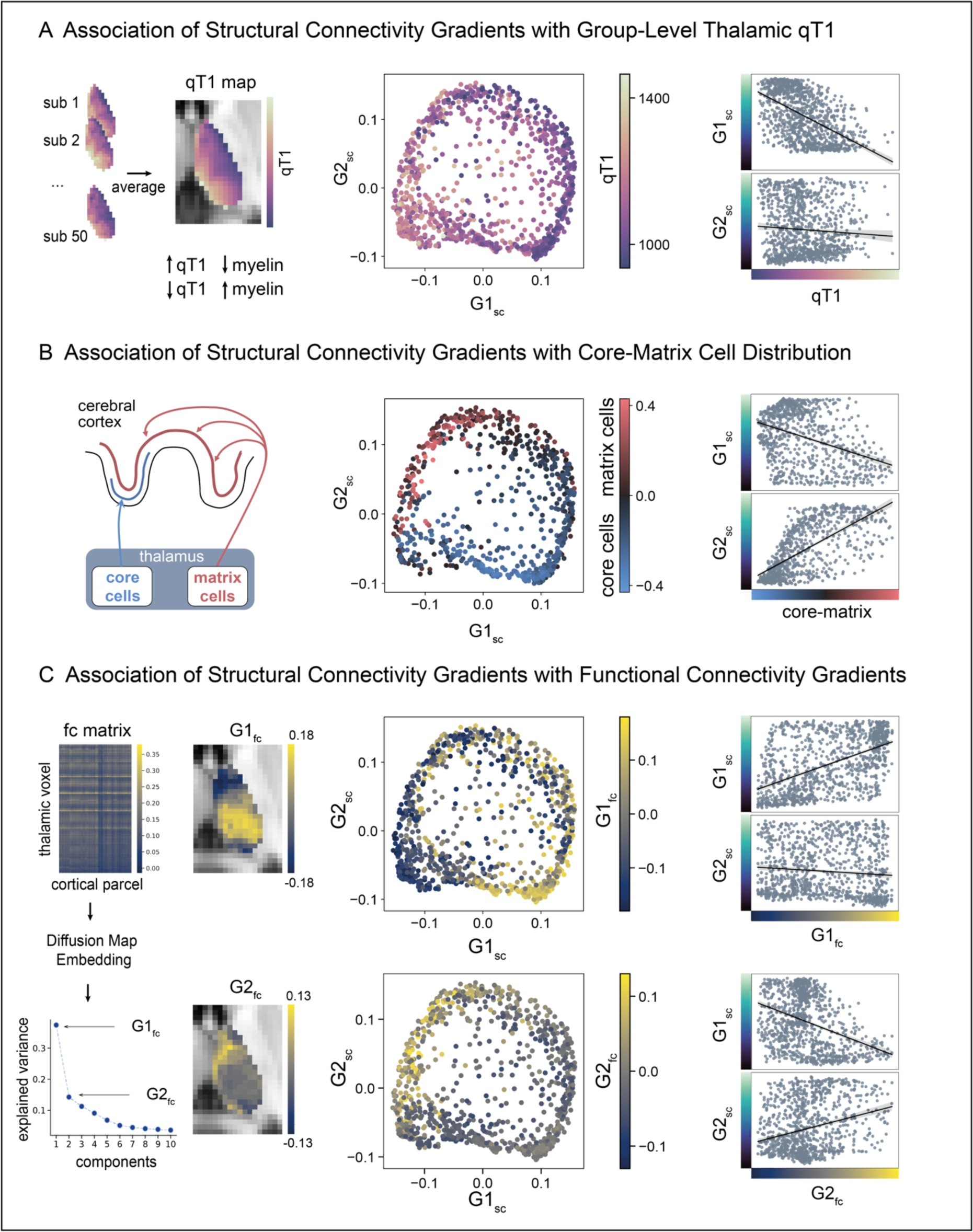
Contextualization of Gradients with Microstructure and Functional Connectivity. A. (left) Individual thalamic qT1 values were averaged to create a group-level qT1 map. Note, the inverse relation between qT1 intensity and approximated grey matter myelin. (middle) 2D space framed by G1_sc_ and G2_sc_, color-coded by thalamic group-level qT1 intensity. (right) Correlation between qT1-intensity and G1_sc_ (upper), and G2 (lower). **B** (left) Conceptualized representation of the core-matrix framework. Core cells (blue) project in a specific fashion to granular layers of the cerebral cortex, whereas matrix cells (red) innervate superficial cortex layers in a distributed fashion. (middle) 2D space framed by G1_sc_ and G2_sc_, color-coded by the core-matrix difference map. Negative values (blue) of the colormap indicate a higher proportion of core cells, whereas positive values (red) indicate a higher proportion of matrix cells. (right) Correlation between core-matrix difference map and G1_sc_ (upper) and G2_sc_ (lower). **C** (left) Functional connectivity matrix (z-scored) resulting from correlating thalamic voxel and cortical parcel time-series and derived gradient decomposition into 10 components with respective eigenvalues. Principal and secondary components (G1_fc_ and G2_fc_) displayed on the axial thalamus slice (see Supplementary Figure 3 for right hemisphere). (middle) 2D space framed by G1_sc_ and G2_sc_, color-coded by (top) G1_fc_ loadings and (bottom) G2_fc_ loadings. (right) Correlation between (top) G1_fc_ and structural connectivity gradients G1_sc_ and G2_sc_, and (bottom) G2_fc_ and structural connectivity gradients G1_sc_ and G2_sc_; All results displayed for the left hemisphere.

#### Intrathalamic Myelin

To examine whether G1_sc_ and G2_sc_ reflect intrathalamic microstructural variation, we used the group-level thalamic qT1 values as a modality estimate for grey matter myelin (Figure 2A). Note that lower qT1 values are associated with a higher myelin content and vice versa. We tested the statistical relationship between both gradient maps and the group-level qT1 map using Pearson correlation and corrected for spatial autocorrelation using variograms (Burt et al., 2020). This analysis suggested a link between G1_sc_ and the spatial distribution of qT1 (LH: r = -0.536, p_SA_ = 0.038), whereas there was no significant correlation between G2_sc_ and the qT1 map (LH: r = -0.068, p_SA_ = 0.873). This trend was replicated in the right thalamus (RH: G1_sc_: r = -0.594, p_SA_ = 0.011; G2_sc_: r = 0.119, p_SA_ = 0.794).

#### Distribution of Core and Matrix Cells

Another main feature of thalamic organization on a microscale level is the varying distribution of core cells and matrix cells that also differ in their TC projection patterns (Jones, 1985; Clascá et al., 2012). To probe whether our identified gradients mirror this distribution, we used a difference map capturing the proportion of core cells and matrix cells based on mRNA level estimates (Müller et al., 2020). Statistically testing the relationship between the core-matrix map and G1_sc_ (LH: r = -0.378, p_SA_ = 0.135) and G2_sc_ (LH: r = 0.676, p_SA_ = 0.044) in the left hemisphere suggested an association between the distribution of core- and matrix cells and G2_sc_ (Figure 2B). This analysis was limited to the left hemisphere due to the small sample size on which the right core-matrix map was grounded (see methods).

#### TC Functional Connectivity Gradients

Next, we explored the link between the low-dimensional organization of structural and functional connectivity (Figure 2C). Therefore, we calculated the TC functional connectivity by correlating the resting-state time series of thalamic seed voxels and cortical parcels and averaged across subjects. Analog to the computation of structural connectivity gradients, we applied diffusion map embedding to uncover the lower dimensional organization of the group-level TC functional connectivity. We correlated the resulting principal and secondary functional connectivity gradient maps (G1_fc_ and G2_fc_) with the structural connectivity gradients and corrected for spatial autocorrelation using variograms. In both hemispheres, G1_fc_ was correlated with G1_sc_ (LH: r = 0.526, p_SA_ = 0.044, RH: r = 0.564, p_SA_ = 0.014). G1_fc_ was not related to G2_sc_ (LH: r = -0.095, p_SA_ = 0.872, RH: r = -0.177, p_SA_ = 0.754). The analysis further did not show an association of G2_fc_ with G1_sc_ (LH: r = -0.374, p_SA_ = 0.083, RH: r = -0.133, p_SA_ = 0.532) and G2_sc_ in the left hemisphere (r = 0.265, p_SA_ = 0.242) but was correlated with G2_sc_ in the right hemisphere (r = 0.484, p_SA_ = 0.016). Taken together, we could show that TC structural connectivity gradients differentially reflected microstructure and cellular properties. Further, we observed an association between TC structural connectivity gradients and intrinsic TC functional connectivity organization.

### Cortical Projections of Structural Connectivity Gradients and Their Associations to Functional Connectivity and Structural Covariance (Figure 3)

Next, we explored the thalamocortical associations based on white matter connectivity, functional connectivity and structural covariance to understand how thalamic and cortical organization interrelate.

**Figure 3:**
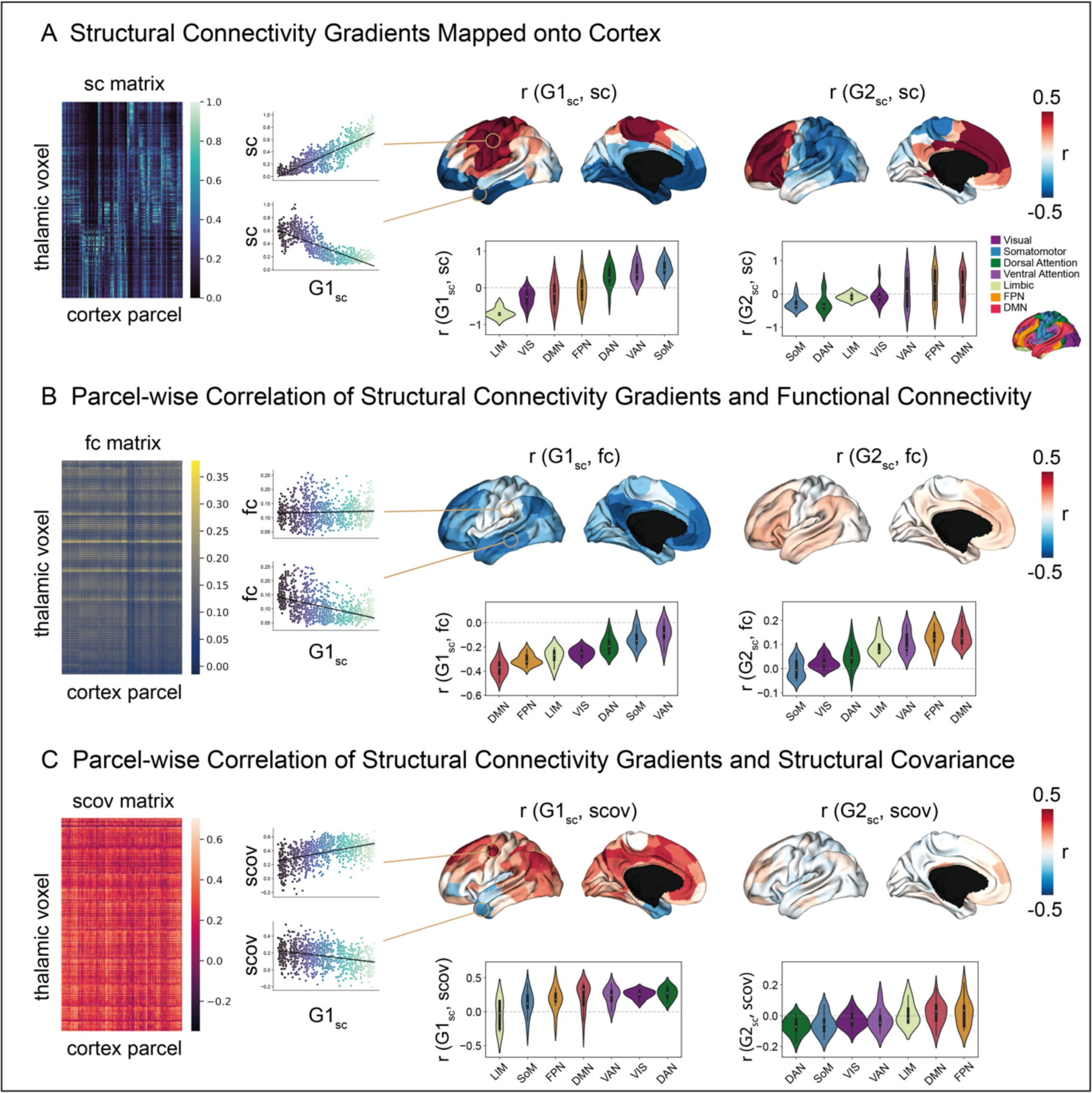
Cortical Projections of Structural Connectivity Gradients and their Associations to Functional Connectivity and Structural Covariance. A. (left) For each cortical parcel, the structural connectivity profiles of the sc matrix were pearson correlated with the structural connectivity gradients. (middle, top) Projection of r values resulting from parcel-wise correlation between sc profiles and G1_sc_ onto cortex. Thus, negative values (blue) indicate relation with the medial part of thalamus, whereas positive values (red) indicate relation with lateral thalamic portions. (middle, bottom) Decoding of cortical pattern leveraging functional communities (ordered along mean). (right) Analog parcel-wise correlation between sc profiles and G2_sc_. **B** (left) Fc matrix and parcel-wise Pearson correlation of resting-state fc strength with the structural connectivity gradients. (middle, top) Projection of r values resulting from parcel-wise correlation between functional connectivity profiles and G1_sc_ onto cortex, and (bottom) decoding of cortical pattern leveraging functional communities (ordered along mean). (right) Analog parcel-wise correlation between functional connectivity profiles and G2_sc_. **C** (left) Scov matrix and parcel-wise Pearson correlation of thalamocortical structural covariance with the structural connectivity gradients. (middle, top) Projection of r values resulting from parcel-wise correlation between structural covariance profiles and G1_sc_ onto cortex, and (bottom) decoding of cortical pattern leveraging functional communities (ordered along mean). (right) Analog parcel-wise correlation between structural covariance profiles and G2_sc_. Abbreviation: sc: structural connectivity, fc: functional connectivity, scov: structural covariance

#### Cortical Projection of TC Structural Connectivity Gradients

First, we computed for each cortical parcel the correlation between the TC structural connectivity profile and G1_sc_. The resulting Pearson’s r value was projected onto the equivalent cortical parcel (Figure 3A). Negatively loaded cortical regions indicated a stronger relationship with the thalamic medial portion, whereas positively loaded regions indicated a stronger relation with the lateral-central subregions of the thalamus. To decode the macroscale cortical patterns, the correlation coefficients were grouped based on functional communities (Yeo et al., 2011) and sorted along the mean r values per community. The decoding revealed a trend following the functional hierarchy from limbic to somatomotor networks with the visual network deviating from this trend. It is worth noting that the lateral geniculate nucleus (LGN, visual projections) is not included in the thalamic mask due to its peripheral localization. Mapping of G2_sc_ onto the cortical surface followed the same principle and resulted in a dissociation between posterior and anterior cortical areas. More specifically, negative loadings located in the somatomotor, and dorsal attention network, whereas positive loadings were situated at the prefrontal cortex and cingulum (Figure 3A). Patterns were reproduced for the right hemisphere (Supplementary Figure 4). Summed up, the principal structural connectivity gradient mapped onto the cortex differentiated somatomotor and limbic areas, while the secondary gradient differentiated between anterior and posterior cortical regions.

#### Associations of TC Structural Connectivity Gradients with TC Functional Connectivity

Next, to study the association between structural connectivity gradients and functional connectivity, G1_sc_ and G2_sc_ were correlated with the functional connectivity profiles of each parcel. The resulting Pearson correlation coefficients were projected onto the cortical surface (Figure 3B). For G1_sc_, negative loadings that indicate a strong relation with the medially located apex of G1_sc_ were present in regions associated with the default mode network (DMN) but also frontoparietal network (FPN), and limbic networks. Positive correlations between parcel-wise functional connectivity profiles and G1_sc_ that would indicate a relation to the more lateral-central thalamic apex were almost not present (LH: 3 % of parcels in range 0 ≤ r ≤ 0.03, RH: 8 % of parcels in range 0 ≤ r ≤ 0.09). For G2_sc_ and parcel-wise functional connectivity profiles, negative correlation coefficients were generally low (LH: 10 % of parcels in range -0.05 ≤ r, RH: 1 % of parcels in range 0 ≤ r). Positive correlation coefficients were present in regions of DMN, FNP, and ventral attention network (VAN). Patterns were reproduced for the right hemisphere (Supplementary Figure 4).

#### Associations of TC Structural Connectivity Gradients with TC Structural Covariance

Last, we aimed to descriptively assess to what extent the defined structural connectivity gradients reflect patterns of TC structural covariance, a global measure that is hypothesized to capture maturational coherence (Alexander-Bloch et al., 2013b). Therefore, we computed a TC structural covariance matrix by correlating the qT1 values of each thalamic voxel with the qT1 value of each cortical parcel (collapsed along cortical depth) across subjects. Next, we correlated G1_sc_ and G2_sc_ with the parcel-wise structural covariance profiles and mapped the results onto the cortical surface (Figure 3C). For G1_sc_, negative correlations indicate stronger covariance between medially located thalamic areas and the temporal pole, lateral frontopolar cortex, and parts of the superior temporal gyrus. Positive loadings showed a dispersed pattern in superior parts of the cerebral cortex, where highest correlations were present in superior frontal, parieto-occipital regions, precuneus, and cingulate gyrus associated with lateral portions of the thalamus. Highest loadings were found in the dorsal attention network (DAN). Here, results are reported for the left hemisphere (Figure 3C). The pattern in the right hemisphere (Supplementary Figure 4) deviated partially (*i.e.*, positive loadings in the superior temporal gyrus), possibly linked to asymmetries in cortical microstructural variability across individuals (Supplementary Figure 5). For G2_sc_ correlation coefficients were generally low (LH: -0.18 ≤ r ≤ 0.21, Figure 3C; RH: -0.28 ≤ r ≤ 0.22; Supplementary Figure 4).

Last, to assess consistency of gradient observations, we selected three prominent example nuclei of the thomas atlas (AV, VPL, and MD) that are distributed along G1_sc_, and explored the structural and functional connectivity projections, and structural covariance patterns. Again, we found the most clear differentiation of thalamic cortical projections for structural connectivity, followed by functional connections. Covariance showed most deviating patterns from the known projection patterns of the nuclei, as well as interhemispheric differences (Supplementary Figure 6).

## 3. Discussion

In this study, we aimed to characterize the thalamic organization based on its structural connectivity profiles to the neocortex beyond the level of distinct nuclei borders. We followed a multimodal approach to explore the association between the defined axes of structural connectivity based thalamic organization and intrathalamic microstructural properties, TC functional connectivity, and TC structural covariance, a proxy for maturational coherence. Our study reported a principal axis/gradient of thalamic organization spanning from medial-to-lateral portions and segregating thalamic higher-order nuclei and sensorimotor nuclei. We found that this pattern spatially correlated with the intrathalamic qT1 distribution, which we used as a proxy for microstructure, and with the first gradient based on TC functional connectivity. Mapped onto the cortex, the principal structural connectivity gradient separated limbic regions and somatomotor regions. Evaluating the multimodal convergence of TC associations (structural connectivity, functional connectivity, and structural covariance), we observed for the principal axis generally a differentiation between limbic, FPN, and DMN regions from DAN and VAN across modalities. Notably, associations with visual and sensorimotor regions were inconsistent, and might arise from modality specific differences or methodological noise. The secondary axis/gradient reflected the spatial distribution of core and matrix cells, and was associated with the secondary gradient based on TC functional connectivity in the right hemisphere. Mapped on the cortex, the secondary gradient differentiated cortical anterior from posterior regions, showed associations between the positive gradient apex and functional connectivity strength in the FPN and DMN, while generally displaying a relatively weak association to structural covariance.

Based on TC white matter connectivity, our findings suggested a principal thalamic medial-to-lateral gradient. Decoding this axis by thalamic nuclei, using a well-established atlas (Saranathan et al., 2021), revealed a trend of medial portions capturing higher-order thalamic nuclei that are known to project to transmodal regions (*i.e.,* AV, MD), while the lateral portions captured nuclei connected to sensorimotor regions (i.e. VLP, CM). This was consistent with the pattern that emerged when the principal axis was projected onto the cortex, which displayed a division between cortical limbic and somatomotor areas. Taken together, we derived that main variations in TC connectivity profiles map an axis differentiating unimodal regions and transmodal regions. Our observations broadly aligned with recent work using a joint analysis of TC structural connectivity and gene expression data (Oldham and Ball, 2023) that reported a similar medial-to-lateral axis, with the difference that in our case the medial apex tended to be more pronounced to the center of the thalamus. The hierarchical representation of projection patterns was also found in the mouse thalamus after computing an organizational axis based on tract tracing and gene expression data, spanning from somatosensory regions to lateral/frontal regions and may suggest the hypothesis that this axis is phylogenetically-conserved (Oldham and Ball, 2023). Further, the medial-to-lateral axis may originate from the spatiotemporal development of the brain. It has been shown that the formation of connectivity-based subdivisions of the human thalamus expanded from lateral to medial portions during the perinatal period (Zheng et al., 2023). Moreover, transcriptional profiling in mice revealed gradual patterns of gene expression that followed this thalamic axis and reflected variations of cellular morphological and electrophysiological properties (Phillips et al., 2019). Next to a medial-to-lateral axis, we observed a second axis of organization with one apex located at the medial-anterior and posterior pole of the thalamus, and the opposite apex intersecting the thalamus from anterior-lateral to central-medial. This secondary gradient mainly segregated MGN, VPL, AV, Pulvinar, CM, and VLP from Hb, MD, MTT, VA, VLa regions on top of the differentiation observed in the first gradient. Reviewing animal studies (Roy et al., 2022), we suggest that the heterogeneity of thalamic organization does exceed distinct nuclei borders and can be described along gradual axes. In sum, our work revealed patterns of thalamic organization along more than one axis, with the principal axis capturing a distinction of transmodal and unimodal differences in projection patterns.

Having established two axes of intrathalamic structural organization using structural connectivity that differentiate thalamic subareas, we further explored how these axes were associated with microstructure, as probed by qT1, a proxy for gray matter myelin. Indeed, we could show that the principal medial-to-lateral axis corresponded to variations of the intrathalamic myelin profile, where lateral and hence thalamic sensorimotor regions showed higher myelination compared to thalamic higher-order regions. This observation was in line with a recent finding suggesting a higher proportion of oligodendrocytes in the lateral portion of the thalamus (Oldham and Ball, 2023). The distribution of myelin could again touch on the spatiotemporal brain development during the perinatal phase. Zheng et al. (2023) reported a lateral-to-medial development of thalamic microstructure, where fiber integrity (measured by fractional anisotropy, fiber density, and diffusivity) in the lateral thalamus seemed to develop faster compared to medial thalamic portion. Further, our finding of higher myelination in lateral thalamic portions, which captured unimodal nuclei, resonated with observations of higher myelination of unimodal regions in the cerebral cortex (Glasser and Van Essen, 2011) and may be related to a higher conduction velocity in sensorimotor regions compared to transmodal regions (Demirtaş et al., 2019; Burt et al., 2018a). Second, in order to further investigate the relation to thalamic microstructure, we leveraged a map that indicates the weighting of core versus matrix cells in thalamic voxels that was created based on mRNA expression levels of the calcium-binding proteins Parvalbumin and Calbindin (Müller et al., 2020). We found the secondary axis, but not the principal axis, being correlated with the cell type distribution. This axis differentiated cortical anterior from posterior regions. Of note, due to the imbalanced donor distribution (6 donors of the left versus 2 of the right hemisphere), we only tested this association in the left hemisphere. Further work will be needed to more precisely map the distribution of core and matrix cells in the human thalamus using a larger sample size.

We further observed a relation between the low dimensional organization of the principal structural and the principal functional TC connectivity gradient, and in one hemisphere, the secondary structural and secondary functional gradient. The functional connectivity pattern in part recapitulated observations of Yang et al. (2020) and together may point to multiple differential axes of organization within the thalamus. While demonstrating that there is an overlap of organizational principles across modalities, differences between patterns of structural and functional connectivity are expected due to method- specificities (*i.e.*, functional connectivity arising not only from direct but also indirect connections).

Indeed, also in parcellation approaches, it has been shown that DWI based clusters correspond higher with structural parcellations in larger nuclei, while parcellations based on resting-state functional MRI (rsfMRI) agree more with structural parcellation in smaller nuclei (Iglehart et al., 2020). Generally, the observation that different organizational dimensions within the thalamus correspond to different structural and functional features indicates that the thalamus and its subnuclei can be differentiated based on different neurobiological principles. It is possible that the divergence observed between the principal and secondary gradient relates to differentiable influences of activity-dependent plasticity and maturation (based on the association of the principal gradient with myelin-proxy profiles), whereas the secondary gradient reflects a different, yet related, organization of core and matrix cells that is scaffolding development but not malleable. However, further work is needed to understand and test the association between cell-level differentiation and possible maturation-related myelination profiles in the thalamus.

Lastly, we aimed to explore how defined patterns of thalamic organization are interrelated with cortical patterns based on structural connectivity, functional connectivity and structural covariance. Evaluating the cortical projections of the principal structural connectivity gradient revealed a differentiation between limbic functional networks that were mostly associated with medial thalamic portions, and the somatomotor network that was related to the lateral thalamic portions. A comparable pattern was observed in functional connectivity. The overall distinction of cortical somatomotor and transmodal projections is in line with the pattern that was revealed by ordering the thalamic nuclei along the principal gradient demonstrating a trend from higher-order nuclei to sensorimotor nuclei. Of note, in the cortical projection patterns, we found the visual network closer at the gradient apex that was in our model assigned to transmodal regions and was therefore deviating from the sensorimotor-association functional hierarchy. This observation may arise due to the exclusion of the LGN from our thalamic mask (projects to primary visual cortex; Jones, 1985; Leh et al., 2008), and further may be driven by the projections of the Pulvinar encompassing higher-order areas and the visual system (Barron et al., 2015). The distinction of sensorimotor and association areas based on structural and functional connectivity aligned with previous work in humans and animals (Jones, 1998; Mukherjee et al., 2020; Harris et al., 2019; Howell et al., 2023) that illustrated how the thalamus may be a key node in the brain coordinating both sensorimotor and abstract cognitive functions (Shine et al., 2023; Wolff et al., 2021). Moreover, the cortical projection pattern of the principle structural connectivity gradient echoed maps of laminar differentiation and sensory-transmodal axes, patterns possibly linked to cortical maturation during the first two decades of human development (Larsen et al., 2023; Sydnor et al., 2023; Burt et al., 2018b; Margulies et al., 2016). Also, this cortical pattern aligned to some extent with notions describing a sensory-fugal gradient of cytoarchitectural complexity (Paquola et al., 2019; Mesulam, 1998), where sensory areas have koniocortical characteristics of a well-developed granular layer IV and fugal or paralimbic areas exhibit a dys/agranular cytoarchitecture. Aggregating our findings and the mentioned literature, the principal gradient derived from TC structural connectivity may in both the thalamus and the cortex be related to development, microstructure and functional hierarchies.

The cortical projection of the secondary structural connectivity gradient revealed a combination of cortical posterior regions (including somatomotor, DAN, limbic, and visual regions) with the negative thalamic apex (medial-anterior and posterior pole of the thalamus) and cortical anterior regions related to the positive thalamic apex (intersection from anterior-lateral to central-medial). Though less pronounced, the pattern was to some extent reflected in TC functional connectivity. The cortical projection of the secondary axis substantially resembled the cortical pattern reported in Müller et al. (2020) that represents the correlation between TC functional connectivity strength and the relative distribution of core and matrix cells, which was consistent with our observation that the secondary structural connectivity gradient is correlated with the relative difference of core and matrix cells in the thalamus. Thalamic regions that contain relatively higher proportions of core cells showed preferential functional coupling to somatosensory cortices, while regions with higher matrix cell proportions preferentially couple with transmodal cortical regions (Müller et al., 2020). Additionally, they found that matrix cell regions tend to couple to cortical areas with a lower intrinsic timescale (Müller et al., 2020). Thus, overall both gradients pointed to a differentiation between sensorimotor and transmodal networks, mirroring observations in recent work of a sensorimotor and association ‘motif’ of thalamic cortical patterning (Howell et al., 2023). Extending this work, we illustrated how these motifs may be embedded within the intrinsic organization of the thalamus along two main structurally defined gradients. Moreover, we found high correspondence between cortical projections of the structural connectivity gradients and functional connectivity patterns.

Throughout the course of development, the thalamus and cortex are closely interconnected. Therefore, we probed whether regions that share similarities in TC connectivity are associated with structural covariance which has been suggested to reflect shared maturation (Alexander-Bloch et al., 2013b). However, this analysis yielded less clear patterns. In the left hemisphere, we observed the medial part of the principal structural connectivity gradient being linked to the temporal pole and hence mostly limbic regions. Lateral thalamic portions were linked to a dispersed pattern of superior regions, while peaking in visual and dorsal attention networks. Noteworthy, the somatomotor network tended to not show a differential association between both anchors of the principal gradient, possibly suggesting more global and unspecific effects of covariance. Further, in contrast to structural and functional connectivity, we found that the pattern of structural covariance deviates between the left and right hemisphere which may be linked to asymmetries in microstructure across individuals. In the cortex, previous studies have shown that there is a strong association between structural covariance and connectivity between areas (Gong et al., 2012; Lerch et al., 2006; Segall et al., 2012). This observation can be associated with the framework of the ‘structural model’, stating that cortical areas with a similar microstructure, in particular laminar differentiation, are also more likely to be structurally and functionally linked (García-Cabezas et al., 2019; Barbas, 2015, 1986; Beul et al., 2017). Extending on studies focusing on structural covariance in the cortex, structural covariance between the thalamus and cortex has been used to parcellate the thalamus in mice (Yee et al., 2024). This work suggested that thalamocortical regions that were connected tend to structurally covary. At the same time, not all structurally covarying regions were connected, possibly pointing to indirect pathways for maturation and connectivity (Yee et al., 2024).

Translating this approach to humans, in the current work, we also found that structural connectivity was not paralleled by structural covariance in all regions. One possibility for explaining structural covariance could be the coactivation of physically connected regions and thereby activity-induced synaptogenesis in a coordinated fashion (Evans, 2013). An alternative explanation might be that structural covariance arises from coordinated developmental processes (Alexander-Bloch et al., 2013b) associated with transcriptomic similarity (Yee et al., 2018). Of note, we found much stronger and differentiable patterns of structural covariance along the principal than the secondary gradient. Adding to the observation of our principal gradient resembling an axis shown to be related to genes that are relevant for thalamic development (Oldham and Ball, 2023) and transcriptional profiling in mice (Phillips et al., 2019), this supported the interpretation of the principal gradient being a more developmentally guided pattern.

## Limitations

To study the organization and connectivity of the human thalamus, the current study was based on *in vivo* MRI. Compared to *post-mortem* studies this comes with the advantage of straightforward data collection of larger sample sizes but with the caveats of noise and lower spatial resolution. In contrast to tracer studies, we note that probabilistic tractography does not reflect connectivity at an axonal level but rather estimates larger fiber tracts and can lead to false positives. Inaccuracies can occur with sharply curved, closely neighboring, or poorly myelinated connections. Although, it has been reported that probabilistic tractography results correspond well to white matter anatomy (Dyrby et al., 2007). Furthermore, we note that in this study only TC connectivity is considered, however, it is well-known that the thalamus is also strongly connected with the subcortex. Notably, the LGN is not included in our thalamic mask due to its extreme posteroventral peripheral location. Furthermore, exploring gradual differences in thalamic organization does not preclude the existence of thalamic subnuclei which can be cytoarchitectonically and functionally delineated with sharp borders. However, we propose that clustering approaches might not account for all overlaps between connectivity profiles in the thalamus and investigating gradual variations could help to understand thalamic organization principles.

In sum, in the current work, we illustrated how the intrinsic organization of the human thalamus, as defined by TC white matter connections derived from DWI, mirrors thalamic microstructure and functional connectivity, and aligns with distributed structural and functional projections shaping cortical organization. In particular, we could identify two axes of thalamic organization, which recapitulate differentiable thalamocortical structural and functional connections and may be rooted in differentiable neurobiological mechanisms of development and maturation. Future work using multimodal high-resolution imaging as well as inclusion of cognitive tasks may help to understand how the anatomy of the thalamus matures and shapes cortical structure, intrinsic function, and ultimately cognitive functional processes.

## 4. Methods

### MRI Data Acquisition

For this study, we used the openly available multimodal MRI dataset for Microstructure-Informed Connectomics (MICA-MICs; Royer et al., 2022). This dataset was acquired from 50 healthy adults (23 women; 29.54 ± 5.62 years) and can be downloaded from the Canadian Open Neuroscience Platform’s data portal (https://portal.conp.ca). The complete cohort underwent a multimodal scanning protocol including high-resolution T1-weighted MRI (T1w), myelin-sensitive qT1 relaxometry, DWI, and rsfMRI at a field strength of 3T. Scanning was conducted at the Brain Imaging Centre of the Montreal Neurological Institute using a 3T Siemens Magnetom Prisma-Fit scanner equipped with a 64-channel head coil. The original study was approved by the ethics committee of the Montreal Neurological Institute and Hospital. Exact acquisition protocols are described elsewhere (Royer et al., 2022). In brief they contained the following:

*T1w:* Using a 3D magnetization-prepared rapid gradient-echo sequence (MP-RAGE), two T1w images with identical parameters (0.8 mm isotropic voxels, matrix = 320 × 320, 224 sagittal slices, TR = 2300 ms, TE = 3.14 ms, TI = 900 ms, flip angle = 9°, iPAT = 2, partial Fourier = 6/8) were acquired.

*DWI*: For the acquisition of the multi-shell DWI data a spin-echo echo-planar imaging sequence was used. This sequence consists of three shells with b-values 300, 700, and 2000s/mm^2^ and 10, 40, and 90 diffusion weighting directions per shell, respectively (1.6 mm isotropic voxels, TR = 3500 ms, TE = 64.40 ms, flip angle = 90°, refocusing flip angle = 180°, FOV = 224 × 224 mm^2^, slice thickness = 1.6 mm, multi-band factor = 3, echo spacing = 0.76 ms, number of b0 images = 3). Additionally, b0 images in reverse phase encoding directions are provided for distortion correction of DWI scans.

*rsFMRI:* Resting-state fMRI images were acquired during a 7 min scan session using multiband accelerated 2D-BOLD echo-planar imaging (3 mm isotropic voxels, TR = 600 ms, TE = 30 ms, flip angle = 52°, FOV = 240 × 240 mm^2^, slice thickness = 3 mm, mb factor = 6, echo spacing = 0.54 ms). For distortion correction of fMRI scans two spin-echo images with reverse phase encoding (3 mm isotropic voxels, TR = 4029 ms, TE = 48 ms, flip angle = 90°, FOV = 240 × 240 mm^2^, slice thickness = 3 mm, echo spacing = 0.54 ms, phase encoding = AP/PA, bandwidth = 2084 Hz/Px) were acquired. Participants HC001 to HC004 underwent slightly longer acquisition (800 time points) but for consistency, we use the same number of time points for all subjects (700 time points).

*qT1:* The qT1 relaxometry data were acquired using a 3D magnetization prepared 2 rapid acquisition gradient echoes sequence (MP2RAGE; 0.8 mm isotropic voxels, 240 sagittal slices, TR = 5000 ms, TE = 2.9 ms, TI 1 = 940 ms, T1 2 = 2830 ms, flip angle 1 = 4°, flip angle 2 = 5°, iPAT = 3, bandwidth = 270 Hz/px, echo spacing = 7.2 ms, partial Fourier = 6/8). To reduce sensitivity to B1 inhomogeneities and to optimize intra- and inter-subject reliability, two inversion images were combined for qT1 mapping.

### Preprocessing

For the preprocessing of MRI data, we used the specific modules of the containerized multimodal MRI processing tool *micapipe* (v. 0.1.2; Cruces et al., 2022) to ensure robustness and reproducibility. Details of the pipeline are described in Cruces et al. (2022).

The below described modality-specific preprocessing steps depend on the output of the module for preprocessing T1w images including reorientation to LPI Orientation, linear alignment and averaging of T1w-scans, intensity nonuniformity correction (N4, Tustison et al., 2010), intensity normalization and a nonlinear registration to MNI152 using ANTS (Tustison and Avants, 2013). Building up on this, micapipe runs FreeSurfers’ recon-all pipeline to segment the cortical surface from native T1w scans (Fischl, 2012).

#### DWI

The used DWI preprocessing module comprises the alignment of multi-shell DWI scans via rigid-body registration, denoising using the Marchenko-Pastur PCA (MP-PCA) algorithm (Veraart et al., 2016), Gibbs ringing correction (Kellner et al., 2016), and the correction of susceptibility-induced geometric distortions, eddy current-induced distortions, and head movements (Andersson et al., 2003; Smith et al., 2004). Further, a non-uniformity bias field correction was performed (Tustison et al., 2010).

#### rsfMRI

The rsfMRI time series data were preprocessed using the adequate micapipe module. The first five volumes were dropped to guarantee magnetic field saturation. Images were reorientated (LPI), motion corrected by registering all time-point volumes to the mean volume, corrected for distortion using the main phase and reverse phase field maps, and denoised using an ICA-FIX classifier (Griffanti et al., 2014; Salimi-Khorshidi et al., 2014). Further, motion spikes were removed using FSL and the average time series was used for registration to FreeSurfer space. The cortical time series were smoothed with a 10 mm Gaussian kernel and nodes are averaged defined by schaefer 200 parcellation scheme. For the time series extraction of thalamic voxels the preprocessed volumes in native space were warped to MNI152 standard space (isotropic resolution 2 mm) using the ANTS transformation parameters provided by micapipe.

#### quantitative T1 (qT1)

QT1 is a proxy for grey matter myelin and provides an index for microstructure. The parameter refers to the T1 relaxation time in milliseconds, which is lower in fatty tissue compared to aqueous tissue (Weiskopf et al., 2021). Accordingly, note that qT1 reflects the amount of grey matter myelin in an inverse relation. To map the individual qT1 scans to MNI152 2 mm standard space, for each subject the transformation between the uni_T1 map with removed background and the MNI152 1 mm reference image was computed using nonlinear registration. Next, the resulting transformation matrix was applied to the individual qT1 scan and data was downscaled to 2 mm isotropic resolution. The thalamic voxel-wise qT1 values were then extracted using a thalamus mask.

The qT1 intensity profiles per cortical parcel are provided by micapipe. Therefore, the module performs a registration from the native qT1 volume to FreeSurfer native space, followed by the construction of 14 equivolumetric surfaces from pial to white-matter boundary (Waehnert et al., 2014). Based on these surfaces, depth-dependent intracortical intensity profiles at each vertex of the native surface mesh are generated and averaged within each of the cortical parcels (schaefer 200 parcellation; Schaefer et al., 2018). For each subject, this resulted in a matrix containing qT1 values and with the dimensions L x N, where L is the number of equivolumetric compartments and N the number of cortical parcels. Note that these equivolumetric compartments do not align with the anatomical cortex layers. For further analysis the pial and grey matter/white matter surface as well as the medial wall were excluded, resulting in a 12 x 200 matrix per subject.

### Thalamic Mask

The binary thalamic mask in MNI152 standard space (2 mm isotropic resolution) was created based on the Harvard-Oxford subcortical atlas (integrated in FSL; Frazier et al., 2005) for each hemisphere, respectively. To incorporate only thalamic voxels in the subsequent analysis and mitigate signal bleeding i.e., from the third ventricle, the thalamic masks underwent a refinement via manual thresholding (left: dilation of 19 %, right: dilation of 20 %).

### TC Structural Connectivity and Gradient Decomposition

#### Voxel-Wise Distribution Estimates of Diffusion Parameters

To estimate the voxel-wise distribution of fiber orientations in the preprocessed diffusion weighted images, we used FSL’s function *bedpostX* (Bayesian Estimation of Diffusion Parameters; Behrens et al., 2003b). The multi-shell extension of the ball- and sticks model was utilized, with a maximum estimate of three fiber orientations within each voxel (Jbabdi et al., 2012).

#### TC Probabilistic Tractography

On a subject level, probabilistic tractography between each voxel in the thalamic seed mask and each parcel of the cortical termination mask (schaefer 200 parcellation; Schaefer et al., 2018; Yeo et al., 2011) was computed using FSL’s *probtrackx2* (probabilistic tracking with crossing fiber). Tractography was computed in the left and right hemispheres independently, in line with previous studies and due to predominant ipsilateral projections (Oldham and Ball, 2023; Iglesias et al., 2018; Behrens et al., 2003a). Each seed voxel was sampled 5000 times with a curvature threshold of 0.2 and step length of 0.5 mm. The path probability maps were corrected for the length of the pathways (--pd). Considering that seed- and target masks are provided in MNI152 standard space, it was additionally required to pass the transformation matrices between native diffusion and MNI standard space. For each subject, we constructed individualized transformations by within-subject registration of the brain extracted b0 image to the native structural scans using FSL’s *FLIRT* (6 degrees of freedom) concatenated to the nonlinear between-subject registration of native structural scans to MNI standard template in 2 mm resolution using a combination of *FLIRT* and *FNIRT* (12 degrees of freedom).

#### TC Structural Connectivity Matrix Computation

Using the output of *probtrackx2*, for both hemispheres a structural connectivity matrix (thalamic seed voxels x cortical parcels) was computed, containing the numbers of streamlines between seed and target per individual. We averaged the individual connectivity matrices across subjects to form a group-level structural connectivity matrix. The group-level structural connectivity matrix was normalized column-wise by dividing all values of each column by the maximum of this column.

#### TC Structural Connectivity Gradients

To uncover the intrathalamic organization based on the TC structural connectome, we employed nonlinear dimensionality reduction. This results in gradient components representing low dimensionally the variation in the connectivity data in a gradual manner. To this end, several analysis steps were performed using the Python toolbox *BrainSpace* (v. 0.1.3; Vos de Wael et al., 2020). The input TC group-level structural connectivity matrix was thresholded at the 75th percentile, in line with previous work (Vos de Wael et al., 2021), and converted into a non-negative squared affinity matrix using a normalized angle similarity kernel. Next, we applied diffusion map embedding, a non-linear dimensionality reduction technique that belongs to the family of graph Laplacians (Coifman and Lafon, 2006) to derive a low-dimensional embedding from the high-dimensional affinity matrix. In this manifold space are nodes with similar connectivity profiles embedded closer together, whereas nodes with distinct connectivity patterns are located further apart. The algorithm is controlled by the *α* parameter which determines the density of sampling points on a manifold (where 0 to 1 = maximal to no influence). Following recommendations based on previous work (Margulies et al., 2016; Paquola et al., 2019; Valk et al., 2022), we set *α* to 0.5 as that will preserve large scale relations between data points in the embedded space and has been suggested to be relatively robust to noise. After assessing the amount of explained variance for each gradient, we mapped the resulting principal (G1_sc_) and secondary gradient (G2_sc_) onto the thalamic mask.

### Contextualization with THOMAS Atlas

To identify the relationship between discrete defined anatomical thalamic subnuclei and TC structural connectivity gradients, we used the parcellation of the THOMAS (Thalamus optimized multi atlas segmentation) atlas in MNI152 space including 11 nuclei (Su et al., 2019; Saranathan et al., 2021; https://doi.org/10.5281/zenodo.5499504). First, we spanned a 2D space framed by G1_sc_ and G2_sc_ and categorized the thalamic voxels according to the nucleus they are part of. Second, for G1_sc_ and G2_sc_, we extracted per nucleus the corresponding gradient loadings and sorted the nuclei in an ascending order based on the median of the corresponding gradient loadings.

### Multimodal Intrathalamic Spatial Organization

#### Intrathalamic qT1 map

By using the thalamic mask, we extracted the thalamic qT1 values at the subject level. Subsequently, we computed the voxel-wise qT1 average across subjects, resulting in a group-level qT1 map of the thalamus.

#### Cell type map based on gene expression levels of Parvalbumin and Calbindin

To approximate the spatial distribution of cell types in the thalamus, we used a relative difference map of core and matrix cells provided by Müller et al., 2020 and publicly available (https://github.com/macshine/corematrix). The map is derived from mRNA level estimates of genes that express the calcium-binding proteins Parvalbumin (PV) and Calbindin (CB_1_) supplied by the Allen Brain Human Brain Atlas (Hawrylycz et al., 2012). PV and CB_1_ have been shown as adequate markers for distinguishing between core- and matrix cells in the thalamus. Considering the sample size on which the data is grounded, we opted to exclusively use the map of the left hemisphere map (6 donors), while excluding the right hemisphere (2 donors). We determined the intersection of this map and our thalamic mask and refer to it as core-matrix map.

#### Functional connectivity gradients

For each subject, we correlated (Pearson) the intrahemispheric timeseries of thalamic voxels and cortical parcels to create a TC functional connectivity matrix per hemisphere (thalamic voxels x cortical parcels). Rows were Fisher z-transformed. By averaging across subjects, the group-level TC functional connectivity matrix was calculated. In line with previous work (Margulies et al., 2016; Valk et al., 2022), this matrix was thresholded at the 90th percentile and used as input to estimate the functional connectivity gradients using diffusion map embedding, analog to gradient computation of structural connectivity described above. G1_fc_ and G2_fc_ were mapped onto the thalamic mask.

#### Association between TC structural connectivity gradients and intrathalamic features

Next, we explored the association between the spatial thalamic organization based on TC structural connectivity and (a) the distribution of thalamic qT1 that is used as a proxy for grey matter myelin, (b) the relative amount of core and matrix cells, and (c) functional connectivity gradients. Therefore, we spanned a 2D space framed by structural connectivity G1_sc_ and G2_sc_ and color-coded the data points based on (a) the group-level qT1 value, and (b) the relative difference between Calbindin and Parvalbumin levels, and (c) the principal and secondary functional connectivity gradient loadings. To further quantify the relationship, we calculated the Pearson correlation of the structural connectivity gradients with (a) the group-level qT1 map, and (b) the cell type map estimating the relative amount of core- and matrix cells, and (c) the principal and secondary functional connectivity gradients. All analyses were corrected for spatial autocorrelation (SA) using the variogram approach implemented in the brainsmash toolbox (Burt et al., 2020).

### Multimodal TC Spatial Organization

#### Mapping TC structural connectivity gradients on the cortex

To project the low dimensional thalamic organization of TC structural connectivity onto the cortex, we calculated the Pearson correlation of the G1_sc_ and G2_sc_ with each column (representing the parcels) of the group-level TC structural connectivity matrix and projected the resulting Pearson correlation coefficient per parcel onto the cortex surface.

#### Mapping the association between TC structural connectivity gradients and functional connectivity on the cortex

To access the link between the thalamic organization based on TC structural connectivity and TC functional connectivity, we calculated the Pearson correlation of the G1_sc_ and G2_sc_ with each column (-representing the parcels) of the group-level TC functional connectivity matrix and projected the resulting Pearson correlation coefficient per parcel onto the cortex surface.

#### TC structural covariance

We further aimed to investigate whether the TC structural connectivity relates to TC structural covariance. First, we averaged the qT1 values of each cortical parcel across compartments. Second, we computed the intrahemispheric TC structural covariance matrix by Pearson correlating qT1 values of thalamic voxels with qT1 values of cortical parcels across subjects. This resulted in a TC structural covariance matrix per hemisphere (M thalamic voxels x N cortical parcels).

#### Mapping the association between TC structural connectivity gradients and TC structural covariance on the cortex

To probe the link between the thalamic organization based on structural connectivity and TC structural covariance, we computed for each parcel the Pearson correlation coefficient by correlating G1_sc_ and G2_sc_ with each column (-representing the parcels) of the group-level TC structural covariance matrix and projected the results onto the cortical surface.

#### Decoding with functional networks

For each modality, the cortical projection patterns were decoded using functional network communities (Visual, Somatomotor, Dorsal Attention, Ventral Attention, Limbic, Frontoparietal and Default-Mode network; Yeo et al., 2011).

### Supplementary Analysis

#### Robustness of structural connectivity gradients computed with different thresholds

To check robustness of the structural connectivity gradient patterns, we computed gradients by thresholding the input group-level structural connectivity matrix at different percentiles (0, 25, 50, 75, 90).

#### Cross-check structural covariance

Supplementary, we calculated the structural covariance between the left thalamus and the cortical parcels of both hemispheres and correlated the resulting structural covariance profiles parcel-wise with the left G1_sc_ and G2_sc_. We did the same analysis steps for the right hemisphere, respectively.

#### Projections based on THOMAS atlas

Using the THOMAS atlas in MNI space, we calculated the mean projections of the nuclei AV, VPL, and MD for structural connections, functional connectivity, and structural covariance.

## Acknowledgements

The authors want to express their gratitude to the various contributors to the MICA-MICs dataset and for openly sharing the dataset with the neuroscientific community. SLV, AJ, AS, BW, and LHS were supported by the Max Planck Society through the Otto Hahn Award. SLV, BCB and AS are furthermore funded by the Helmholtz International BigBrain Analytics and Learning Laboratory (HIBALL), supported by the Helmholtz Association’s Initiative and Networking Fund and the Healthy Brains, Healthy Lives initiative at McGill University. BCB acknowledges research support from the National Science and Engineering Research Council of Canada (NSERC Discovery-1304413), Canadian Institutes of Health Research (FDN-154298, PJT-174995), SickKids Foundation (NI17-039), BrainCanada, FRQ-S, and the Tier-2 Canada Research Chairs program. JR was supported by a fellowship from the Canadian Institutes of Health Research. MDH was funded by the German Federal Ministry of Education and Research (BMBF) and the Max Planck Society.

## 5. Code and Data Availability

The data used in this study is openly available and can be downloaded from: https://portal.conp.ca/dataset?id=projects/mica-mics. For data preprocessing we used the openly available processing pipeline micapipe (v. 0.1.2; https://micapipe.readthedocs.io/) and FSL (https://fsl.fmrib.ox.ac.uk/fsl/fslwiki). The template of the THOMAS atlas can be requested here: https://doi.org/10.5281/zenodo.5499504. The core-matrix difference is available at: https://github.com/macshine/corematrix. Custom code generated for analysis and plotting is available at a Github repository (https://github.com/CNG-LAB/cngopen/tree/main/thalamic_gradients). Further, our code includes open software packages: The gradient computation was carried out using BrainSpace (v. 0.1.3; https://brainspace.readthedocs.io/en/latest/). For spatial autocorrelation correction, the brainsmash toolbox was used (v. 0.11.0; https://brainsmash.readthedocs.io/en/latest/).

---

**Supplementary Figure 1:**
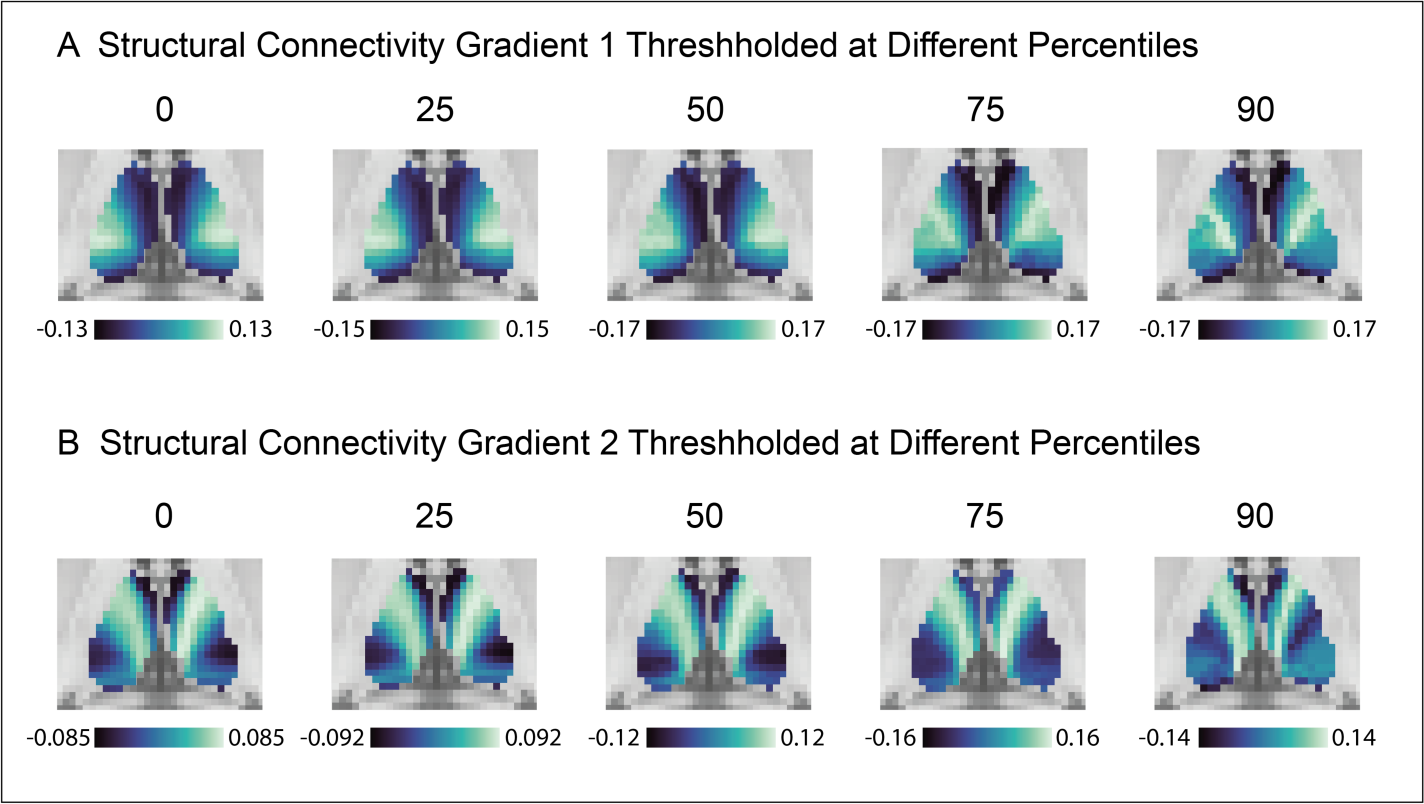
Robustness of Structural Connectivity Gradients. Gradients were derived from the group-level thalamocortical structural connectivity matrix thresholded at different percentiles (0, 25, 50, 75, 90). **A** Gradient loadings of component 1 (G1_sc_) were projected on the thalamus (axial plane). **B** Gradient loadings of component 2 (G2_sc_) were projected on the thalamus (axial plane).

**Supplementary Figure 2:**
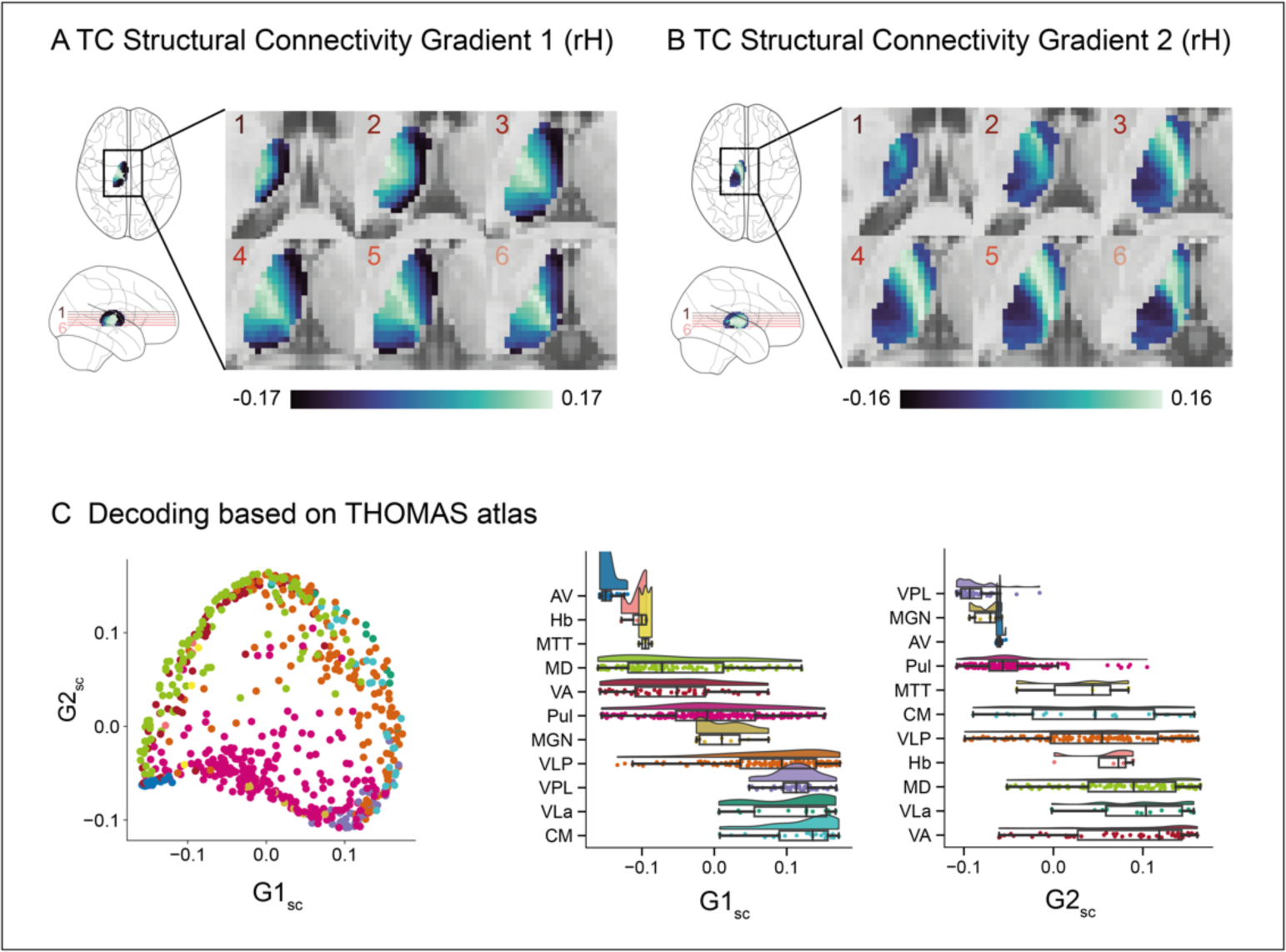
Thalamocortical Structural Connectivity Gradients (RH). A. Gradient loadings of component 1 (G1_sc_) projected on the thalamus (axial planes). The red lines in the glass brain indicate the position of each respective axial slice of the displayed thalamus. **B** Gradient loadings of component 2 (G2_sc_) projected on the thalamus. Slice positions are congruent to A. **C (**left) Decoding of G1_sc_ and G2_sc_ based on THOMAS atlas. 2D space framed by G1_sc_ and G2_sc_, color-coded by thalamic subnuclei. (middle and right) Raincloud plots display the gradient loadings of G1_sc_ and G2_sc_ per nucleus and are ordered by median, respectively. Abbreviations in **C**: AV: Anterior ventral nucleus, VA: Ventral anterior nucleus, VLa: Ventral lateral anterior nucleus, VLP: Ventral lateral posterior nucleus, VPL: Ventral posterior lateral nucleus, Pul: Pulvinar nucleus, MGN: Medial geniculate nucleus, CM: Centromedian nucleus, MD: Mediodorsal nucleus, Hb: Habenular nucleus, MTT: Mammillothalamic tract

**Supplementary Figure 3:**
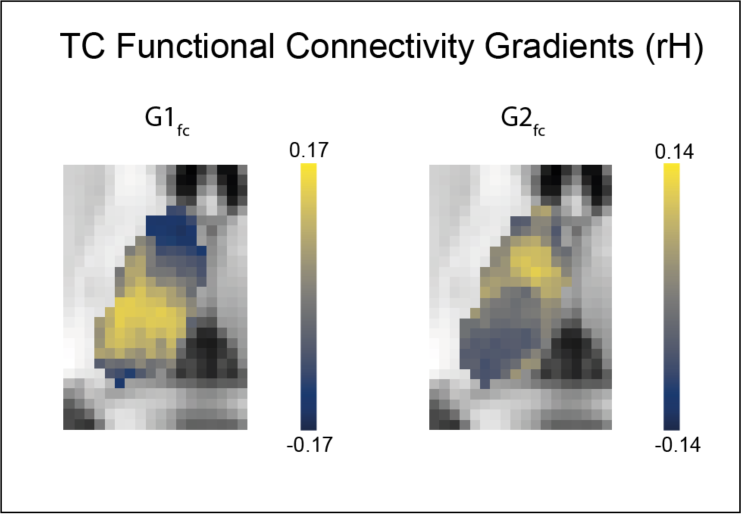
Thalamocortical Functional Connectivity Gradients (RH). Right hemisphere gradient loadings of component 1 (G1_sc_) and component 2 (G2_sc_) projected on the thalamus (axial slice).

**Supplementary Figure 4:**
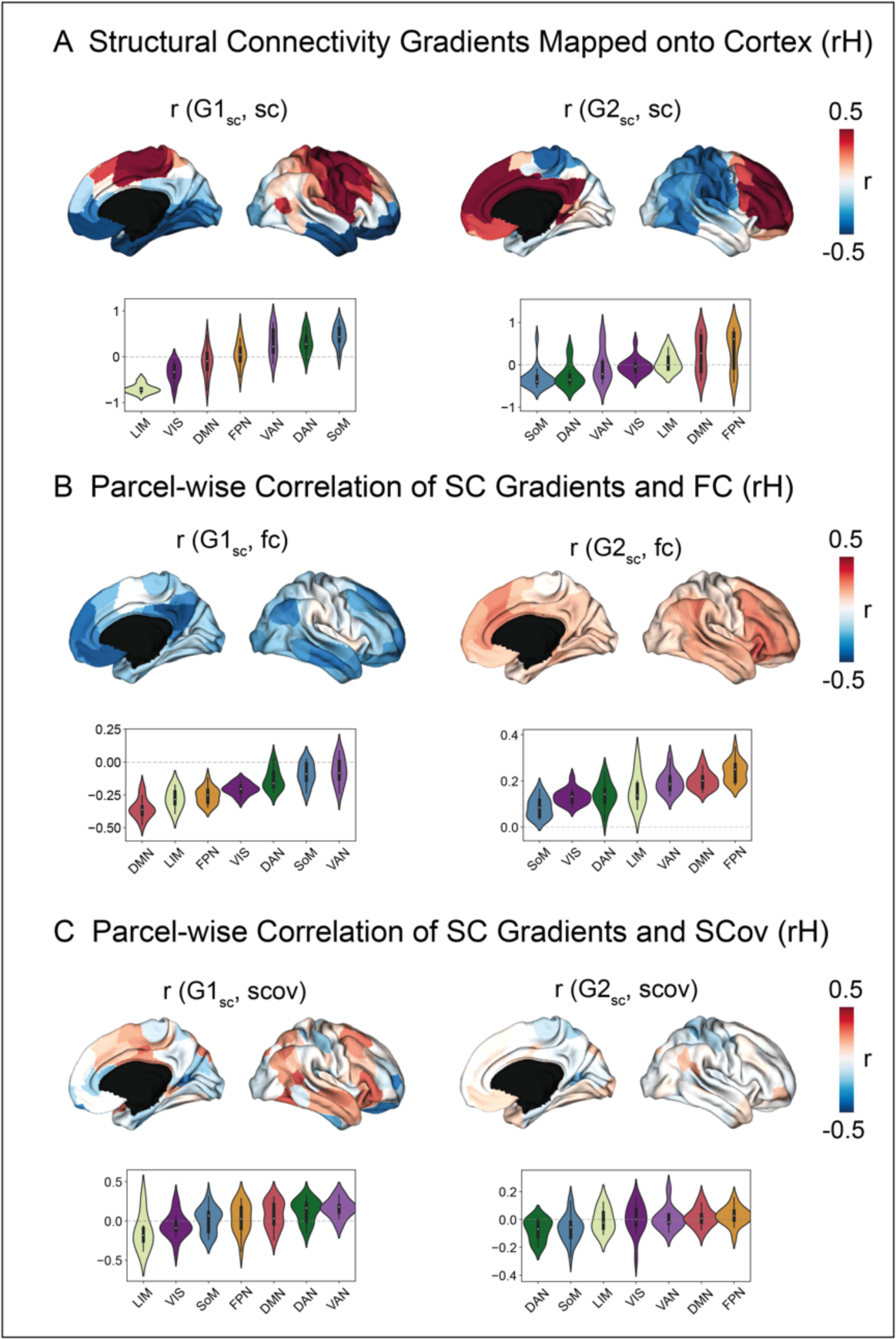
Cortical Projections of Structural Connectivity Gradients and their Association to Functional Connectivity and Structural Covariance (RH). A. (left, top) Projection of r values resulting from parcel-wise correlation between sc profiles and G1_sc_ onto cortex. Thus, negative values (blue) indicate relation with the medial part of thalamus, whereas positive values (red) indicate relation with lateral thalamic portions. (left, bottom) Decoding of cortical pattern leveraging functional communities (ordered along mean). (right) Analog parcel-wise correlation between sc profiles and G2_sc_, and decoding. **B** (left, top) Projection of r values resulting from parcel-wise correlation between functional connectivity profiles and G1_sc_ onto cortex, and (left, bottom) decoding of cortical pattern leveraging functional communities (ordered along mean). (right) Analog parcel-wise correlation between functional connectivity profiles and G2_sc_, and decoding. **C** (left, top) Projection of r values resulting from parcel-wise correlation between structural covariance profiles and G1_sc_ onto cortex, and (left, bottom) decoding of cortical pattern leveraging functional communities (ordered along mean). (right) Analog parcel-wise correlation between structural covariance profiles and G2_sc_, and decoding.

**Supplementary Figure 5:**
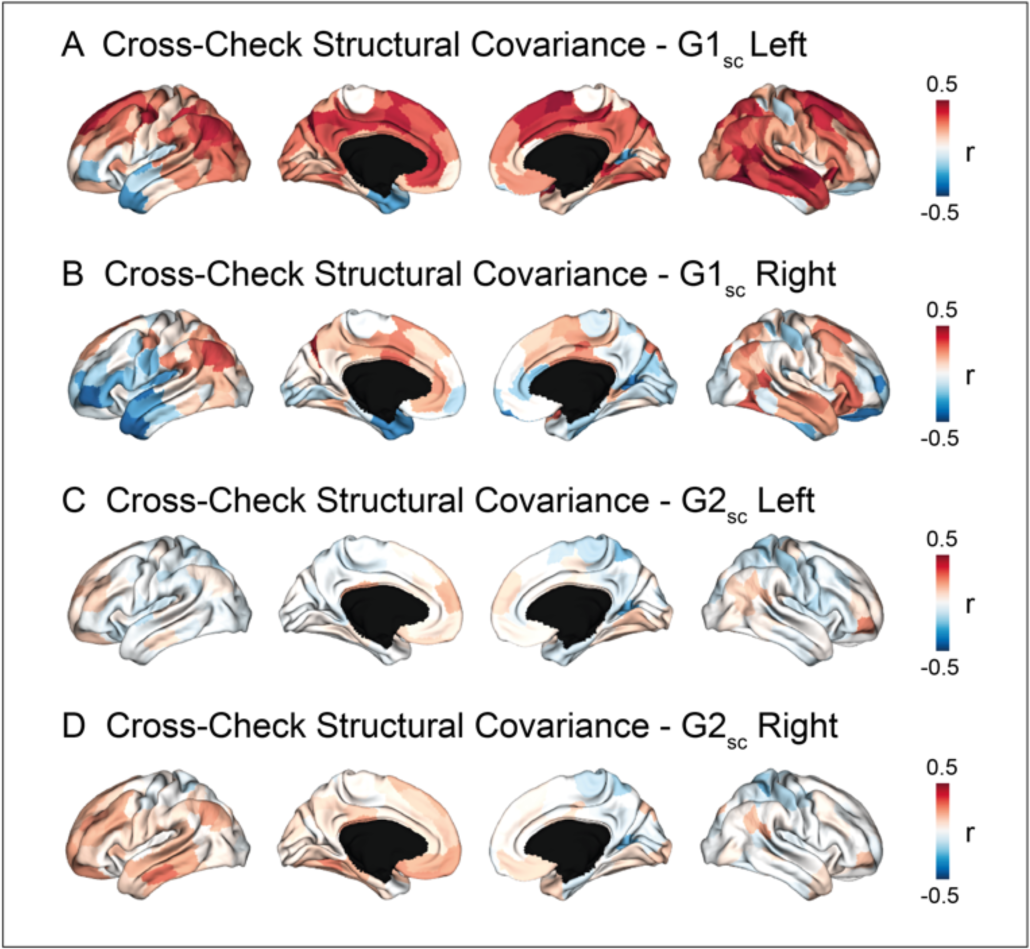
Cross-Check of Structural Covariance Results. A. Structural covariance was computed between left thalamic voxels to both cortex hemispheres. The resulting structural covariance profiles were parcel-wise correlated with the left G1_sc_ and r values were projected onto the cortex. **B** Structural covariance was computed between right thalamic voxels to both cortex hemispheres. The resulting structural covariance profiles were parcel-wise correlated with the right G1_sc_ and r values were projected onto the cortex. **C** Procedure analog to A but with G2_sc_. **D** Procedure analog to B but with G2_sc_.

**Supplementary Figure 6:**
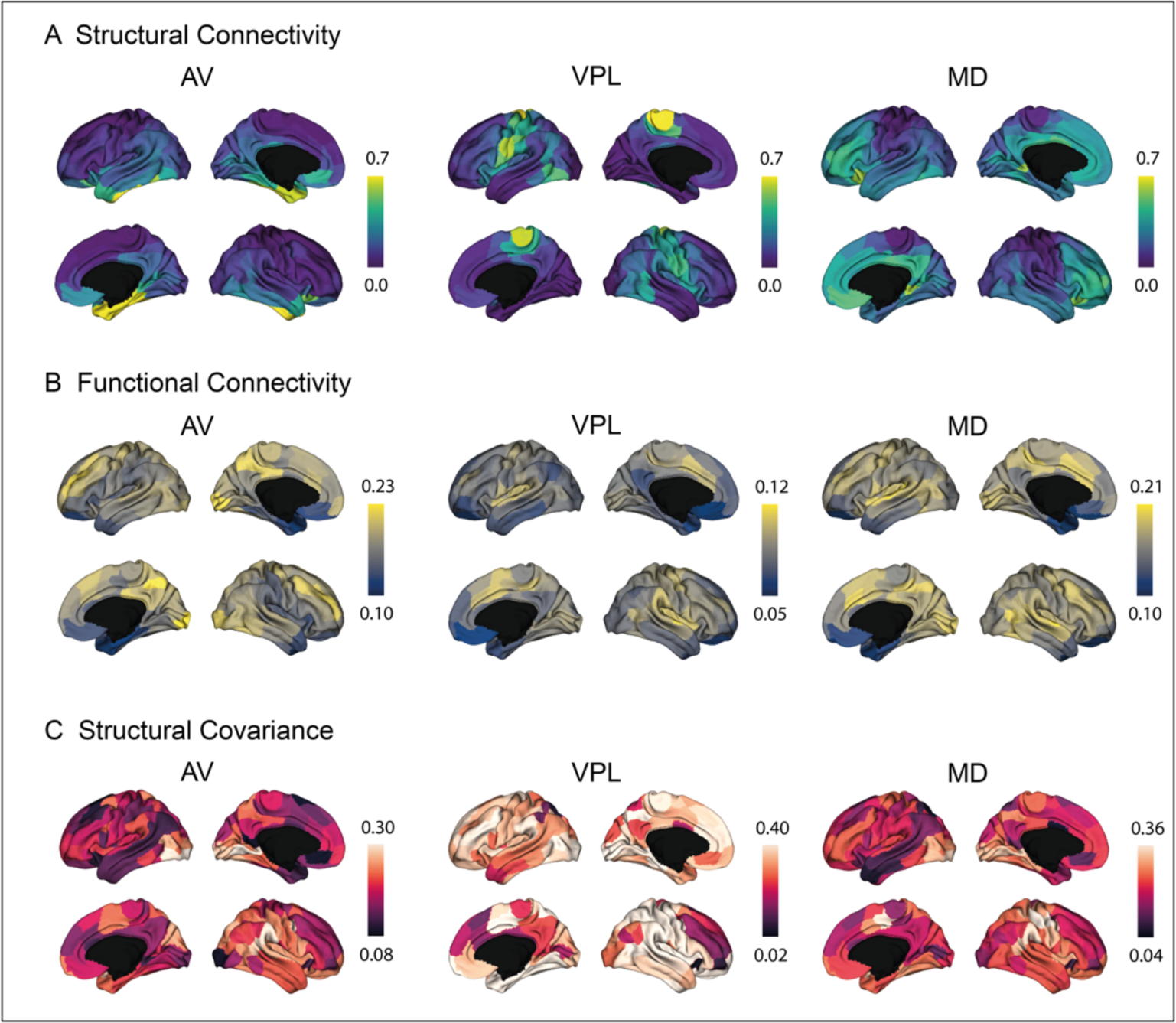
Projections Based on Thomas Nuclei. A. Mean structural connectivity projections from AV, VPL, and MD projected onto the cortex. **B** Mean functional connectivity projections from AV, VPL, and MD projected onto the cortex. **C** Mean structural covariance projections from AV, VPL, and MD projected onto the cortex.

## Footnotes

1 AV: Anterior ventral nucleus, MTT: Mammillothalamic tract, Hb: Habenular nucleus, MD: Mediodorsal nucleus, VA: Ventral anterior nucleus, Pul: Pulvinar nucleus, MGN: Medial geniculate nucleus, VLa: Ventral lateral anterior nucleus, VPL: Ventral posterior lateral nucleus, VLP: Ventral lateral posterior nucleus, CM: Centromedian nucleus (Su et al., 2019)

## References

Alexander-Bloch, Giedd, J.N., Bullmore, E., 2013a. Imaging structural co-variance between human brain regions. Nat Rev Neurosci 14, 322–336. 10.1038/nrn3465

Alexander-Bloch, Raznahan, A., Bullmore, E., Giedd, J., 2013b. The Convergence of Maturational Change and Structural Covariance in Human Cortical Networks. Journal of Neuroscience 33, 2889–2899. 10.1523/JNEUROSCI.3554-12.2013

Andersson, J.L.R., Skare, S., Ashburner, J., 2003. How to correct susceptibility distortions in spin-echo echo-planar images: application to diffusion tensor imaging. NeuroImage 20, 870–888. 10.1016/S1053-8119(03)00336-7

Antón-Bolaños, N., Espinosa, A., López-Bendito, G., 2018. Developmental interactions between thalamus and cortex: a true love reciprocal story. Curr Opin Neurobiol 52, 33–41. 10.1016/j.conb.2018.04.018

Barbas, H., 2015. General Cortical and Special Prefrontal Connections: Principles from Structure to Function. Annu. Rev. Neurosci. 38, 269–289. 10.1146/annurev-neuro-071714-033936

Barbas, H., 1986. Pattern in the laminar origin of corticocortical connections. J. Comp. Neurol. 252, 415–422. 10.1002/cne.902520310

Barron, D.S., Eickhoff, S.B., Clos, M., Fox, P.T., 2015. Human pulvinar functional organization and connectivity. Hum Brain Mapp 36, 2417–2431. 10.1002/hbm.22781

Battistella, G., Najdenovska, E., Maeder, P., Ghazaleh, N., Daducci, A., Thiran, J.-P., Jacquemont, S., Tuleasca, C., Levivier, M., Bach Cuadra, M., Fornari, E., 2017. Robust thalamic nuclei segmentation method based on local diffusion magnetic resonance properties. Brain Struct Funct 222, 2203–2216. 10.1007/s00429-016-1336-4

Behrens, T.E.J., Johansen-Berg, H., Woolrich, M.W., Smith, S.M., Wheeler-Kingshott, C.A.M., Boulby, P.A., Barker, G.J., Sillery, E.L., Sheehan, K., Ciccarelli, O., Thompson, A.J., Brady, J.M., Matthews, P.M., 2003a. Non-invasive mapping of connections between human thalamus and cortex using diffusion imaging. Nat Neurosci 6, 750–757. 10.1038/nn1075

Behrens, T.E.J., Woolrich, M.W., Jenkinson, M., Johansen-Berg, H., Nunes, R.G., Clare, S., Matthews, P.M., Brady, J.M., Smith, S.M., 2003b. Characterization and propagation of uncertainty in diffusion-weighted MR imaging. Magn. Reson. Med. 50, 1077–1088. 10.1002/mrm.10609

Bernhardt, B.C., Smallwood, J., Keilholz, S., Margulies, D.S., 2022. Gradients in brain organization. NeuroImage 251, 118987. 10.1016/j.neuroimage.2022.118987

Beul, S.F., Barbas, H., Hilgetag, C.C., 2017. A Predictive Structural Model of the Primate Connectome. Sci Rep 7, 43176. 10.1038/srep43176

Burt, J.B., Demirtaş, M., Eckner, W.J., Navejar, N.M., Ji, J.L., Martin, W.J., Bernacchia, A., Anticevic, A., Murray, J.D., 2018a. Hierarchy of transcriptomic specialization across human cortex captured by structural neuroimaging topography. Nat Neurosci 21, 1251–1259. 10.1038/s41593-018-0195-0

Burt, J.B., Demirtaş, M., Eckner, W.J., Navejar, N.M., Ji, J.L., Martin, W.J., Bernacchia, A., Anticevic, A., Murray, J.D., 2018b. Hierarchy of transcriptomic specialization across human cortex captured by structural neuroimaging topography. Nat Neurosci 21, 1251–1259. 10.1038/s41593-018-0195-0

Burt, J.B., Helmer, M., Shinn, M., Anticevic, A., Murray, J.D., 2020. Generative modeling of brain maps with spatial autocorrelation. NeuroImage 220, 117038. 10.1016/j.neuroimage.2020.117038

Clascá, F., Rubio-Garrido, P., Jabaudon, D., 2012. Unveiling the diversity of thalamocortical neuron subtypes: Thalamocortical neuron diversity. European Journal of Neuroscience 35, 1524– 1532. 10.1111/j.1460-9568.2012.08033.x

Coifman, R.R., Lafon, S., 2006. Diffusion maps. Applied and Computational Harmonic Analysis 21, 5–30. 10.1016/j.acha.2006.04.006

Cruces, R.R., Royer, J., Herholz, P., Larivière, S., Vos de Wael, R., Paquola, C., Benkarim, O., Park, B., Degré-Pelletier, J., Nelson, M.C., DeKraker, J., Leppert, I.R., Tardif, C., Poline, J.-B., Concha, L., Bernhardt, B.C., 2022. Micapipe: A pipeline for multimodal neuroimaging and connectome analysis. NeuroImage 263, 119612. 10.1016/j.neuroimage.2022.119612

de Bourbon-Teles, J., Bentley, P., Koshino, S., Shah, K., Dutta, A., Malhotra, P., Egner, T., Husain, M., Soto, D., 2014. Thalamic Control of Human Attention Driven by Memory and Learning. Current Biology 24, 993–999. 10.1016/j.cub.2014.03.024

Demirtaş, M., Burt, J.B., Helmer, M., Ji, J.L., Adkinson, B.D., Glasser, M.F., Van Essen, D.C., Sotiropoulos, S.N., Anticevic, A., Murray, J.D., 2019. Hierarchical Heterogeneity across Human Cortex Shapes Large-Scale Neural Dynamics. Neuron 101, 1181–1194.e13. 10.1016/j.neuron.2019.01.017

Dyrby, T.B., Søgaard, L.V., Parker, G.J., Alexander, D.C., Lind, N.M., Baaré, W.F.C., Hay-Schmidt, A., Eriksen, N., Pakkenberg, B., Paulson, O.B., Jelsing, J., 2007. Validation of in vitro probabilistic tractography. NeuroImage 37, 1267–1277. 10.1016/j.neuroimage.2007.06.022

Evans, A.C., 2013. Networks of anatomical covariance. NeuroImage 80, 489–504. 10.1016/j.neuroimage.2013.05.054

Fischl, B., 2012. FreeSurfer. NeuroImage, 20 YEARS OF fMRI 62, 774–781. 10.1016/j.neuroimage.2012.01.021

Frazier, J.A., Chiu, S., Breeze, J.L., Makris, N., Lange, N., Kennedy, D.N., Herbert, M.R., Bent, E.K., Koneru, V.K., Dieterich, M.E., Hodge, S.M., Rauch, S.L., Grant, P.E., Cohen, B.M., Seidman, L.J., Caviness, V.S., Biederman, J., 2005. Structural Brain Magnetic Resonance Imaging of Limbic and Thalamic Volumes in Pediatric Bipolar Disorder. AJP 162, 1256– 1265. 10.1176/appi.ajp.162.7.1256

Gao, C., Leng, Y., Ma, J., Rooke, V., Rodriguez-Gonzalez, S., Ramakrishnan, C., Deisseroth, K., Penzo, M.A., 2020. Two genetically, anatomically and functionally distinct cell types segregate across anteroposterior axis of paraventricular thalamus. Nat Neurosci 23, 217–228. 10.1038/s41593-019-0572-3

García-Cabezas, M.Á., Zikopoulos, B., Barbas, H., 2019. The Structural Model: a theory linking connections, plasticity, pathology, development and evolution of the cerebral cortex. Brain Struct Funct 224, 985–1008. 10.1007/s00429-019-01841-9

Glasser, M.F., Van Essen, D.C., 2011. Mapping Human Cortical Areas In Vivo Based on Myelin Content as Revealed by T1- and T2-Weighted MRI. J Neurosci 31, 11597–11616. 10.1523/JNEUROSCI.2180-11.2011

Gong, G., He, Y., Chen, Z.J., Evans, A.C., 2012. Convergence and divergence of thickness correlations with diffusion connections across the human cerebral cortex. NeuroImage 59, 1239–1248. 10.1016/j.neuroimage.2011.08.017

Govek, K.W., Chen, S., Sgourdou, P., Yao, Y., Woodhouse, S., Chen, T., Fuccillo, M.V., Epstein, D.J., Camara, P.G., 2022. Developmental trajectories of thalamic progenitors revealed by single-cell transcriptome profiling and Shh perturbation. Cell Reports 41, 111768. 10.1016/j.celrep.2022.111768

Griffanti, L., Salimi-Khorshidi, G., Beckmann, C.F., Auerbach, E.J., Douaud, G., Sexton, C.E., Zsoldos, E., Ebmeier, K.P., Filippini, N., Mackay, C.E., Moeller, S., Xu, J., Yacoub, E., Baselli, G., Ugurbil, K., Miller, K.L., Smith, S.M., 2014. ICA-based artefact removal and accelerated fMRI acquisition for improved resting state network imaging. NeuroImage 95, 232–247. 10.1016/j.neuroimage.2014.03.034

Halassa, M.M., Kastner, S., 2017. Thalamic functions in distributed cognitive control. Nat Neurosci 20, 1669–1679. 10.1038/s41593-017-0020-1

Harris, J.A., Mihalas, S., Hirokawa, K.E., Whitesell, J.D., Choi, H., Bernard, A., Bohn, P., Caldejon, S., Casal, L., Cho, A., Feiner, A., Feng, D., Gaudreault, N., Gerfen, C.R., Graddis, N., Groblewski, P.A., Henry, A.M., Ho, A., Howard, R., Knox, J.E., Kuan, L., Kuang, X., Lecoq, J., Lesnar, P., Li, Y., Luviano, J., McConoughey, S., Mortrud, M.T., Naeemi, M., Ng, L., Oh, S.W., Ouellette, B., Shen, E., Sorensen, S.A., Wakeman, W., Wang, Q., Wang, Y., Williford, A., Phillips, J.W., Jones, A.R., Koch, C., Zeng, H., 2019. Hierarchical organization of cortical and thalamic connectivity. Nature 575, 195–202. 10.1038/s41586-019-1716-z

Hawrylycz, M.J., Lein, E.S., Guillozet-Bongaarts, A.L., Shen, E.H., Ng, L., Miller, J.A., van de Lagemaat, L.N., Smith, K.A., Ebbert, A., Riley, Z.L., Abajian, C., Beckmann, C.F., Bernard, A., Bertagnolli, D., Boe, A.F., Cartagena, P.M., Chakravarty, M.M., Chapin, M., Chong, J., Dalley, R.A., Daly, B.D., Dang, C., Datta, S., Dee, N., Dolbeare, T.A., Faber, V., Feng, D., Fowler, D.R., Goldy, J., Gregor, B.W., Haradon, Z., Haynor, D.R., Hohmann, J.G., Horvath, S., Howard, R.E., Jeromin, A., Jochim, J.M., Kinnunen, M., Lau, C., Lazarz, E.T., Lee, C., Lemon, T.A., Li, L., Li, Y., Morris, J.A., Overly, C.C., Parker, P.D., Parry, S.E., Reding, M., Royall, J.J., Schulkin, J., Sequeira, P.A., Slaughterbeck, C.R., Smith, S.C., Sodt, A.J., Sunkin, S.M., Swanson, B.E., Vawter, M.P., Williams, D., Wohnoutka, P., Zielke, H.R., Geschwind, D.H., Hof, P.R., Smith, S.M., Koch, C., Grant, S.G.N., Jones, A.R., 2012. An anatomically comprehensive atlas of the adult human brain transcriptome. Nature 489, 391–399. 10.1038/nature11405

Howell, A.M., Warrington, S., Fonteneau, C., Cho, Y.T., Sotiropoulos, S.N., Murray, J.D., Anticevic, A., 2023. The spatial extent of anatomical connections within the thalamus varies across the cortical hierarchy in humans and macaques. bioRxiv 2023.07.22.550168. 10.1101/2023.07.22.550168

Huntenburg, J.M., Bazin, P.-L., Margulies, D.S., 2018. Large-Scale Gradients in Human Cortical Organization. Trends in Cognitive Sciences 22, 21–31. 10.1016/j.tics.2017.11.002

Hwang, K., Bertolero, M.A., Liu, W.B., D’Esposito, M., 2017. The Human Thalamus Is an Integrative Hub for Functional Brain Networks. J. Neurosci. 37, 5594–5607. 10.1523/JNEUROSCI.0067-17.2017

Hwang, K., Shine, J.M., Cole, M.W., Sorenson, E., 2022. Thalamocortical contributions to cognitive task activity. eLife 11, e81282. 10.7554/eLife.81282

Iglehart, C., Monti, M., Cain, J., Tourdias, T., Saranathan, M., 2020. A systematic comparison of structural-, structural connectivity-, and functional connectivity-based thalamus parcellation techniques. Brain Struct Funct 225, 1631–1642. 10.1007/s00429-020-02085-8

Iglesias, J.E., Insausti, R., Lerma-Usabiaga, G., Bocchetta, M., Van Leemput, K., Greve, D.N., van der Kouwe, A., Fischl, B., Caballero-Gaudes, C., Paz-Alonso, P.M., 2018. A probabilistic atlas of the human thalamic nuclei combining ex vivo MRI and histology. NeuroImage 183, 314–326. 10.1016/j.neuroimage.2018.08.012

Jbabdi, S., Sotiropoulos, S.N., Savio, A.M., Graña, M., Behrens, T.E.J., 2012. Model-based analysis of multishell diffusion MR data for tractography: How to get over fitting problems. Magn Reson Med 68, 1846–1855. 10.1002/mrm.24204

Ji, B., Li, Z., Li, K., Li, L., Langley, J., Shen, H., Nie, S., Zhang, R., Hu, X., 2016. Dynamic thalamus parcellation from resting-state fMRI data. Human Brain Mapping 37, 954–967. 10.1002/hbm.23079

Johansen-Berg, H., Behrens, T.E.J., Sillery, E., Ciccarelli, O., Thompson, A.J., Smith, S.M., Matthews, P.M., 2005. Functional–Anatomical Validation and Individual Variation of Diffusion Tractography-based Segmentation of the Human Thalamus. Cerebral Cortex 15, 31–39. 10.1093/cercor/bhh105

Jones, E.G., 2009. Synchrony in the Interconnected Circuitry of the Thalamus and Cerebral Cortex. Annals of the New York Academy of Sciences 1157, 10–23. 10.1111/j.1749-6632.2009.04534.x

Jones, E.G., 2001. The thalamic matrix and thalamocortical synchrony. Trends in Neurosciences 24, 595–601. 10.1016/S0166-2236(00)01922-6

Jones, E.G., 1998. Viewpoint: the core and matrix of thalamic organization. Neuroscience 85, 331–345. 10.1016/S0306-4522(97)00581-2

Jones, E.G. (Ed.), 1985. The Thalamus. Springer US, Boston, MA. 10.1007/978-1-4615-1749-8

Kellner, E., Dhital, B., Kiselev, V.G., Reisert, M., 2016. Gibbs-ringing artifact removal based on local subvoxel-shifts: Gibbs-Ringing Artifact Removal. Magn. Reson. Med. 76, 1574–1581. 10.1002/mrm.26054

Larsen, B., Sydnor, V.J., Keller, A.S., Yeo, B.T.T., Satterthwaite, T.D., 2023. A critical period plasticity framework for the sensorimotor-association axis of cortical neurodevelopment. Trends Neurosci 46, 847–862. 10.1016/j.tins.2023.07.007

Leh, S.E., Chakravarty, M.M., Ptito, A., 2008. The Connectivity of the Human Pulvinar: A Diffusion Tensor Imaging Tractography Study. Int J Biomed Imaging 2008, 789539. 10.1155/2008/789539

Lerch, J.P., Worsley, K., Shaw, W.P., Greenstein, D.K., Lenroot, R.K., Giedd, J., Evans, A.C., 2006. Mapping anatomical correlations across cerebral cortex (MACACC) using cortical thickness from MRI. NeuroImage 31, 993–1003. 10.1016/j.neuroimage.2006.01.042

López-Bendito, G., 2018. Development of the Thalamocortical Interactions: Past, Present and Future. Neuroscience 385, 67–74. 10.1016/j.neuroscience.2018.06.020

López-Bendito, G., Molnár, Z., 2003. Thalamocortical development: how are we going to get there? Nat Rev Neurosci 4, 276–289. 10.1038/nrn1075

Margulies, D.S., Ghosh, S.S., Goulas, A., Falkiewicz, M., Huntenburg, J.M., Langs, G., Bezgin, G., Eickhoff, S.B., Castellanos, F.X., Petrides, M., Jefferies, E., Smallwood, J., 2016. Situating the default-mode network along a principal gradient of macroscale cortical organization. Proc. Natl. Acad. Sci. U.S.A. 113, 12574–12579. 10.1073/pnas.1608282113

Mesulam, M., 1998. From sensation to cognition. Brain 121, 1013–1052. 10.1093/brain/121.6.1013

Morel, A., Magnin, M., Jeanmonod, D., 1997. Multiarchitectonic and stereotactic atlas of the human thalamus. J. Comp. Neurol. 387, 588–630. 10.1002/(SICI)1096-9861(19971103)387:4<588::AID-CNE8>3.0.CO;2-Z

Mukherjee, A., Bajwa, N., Lam, N.H., Porrero, C., Clasca, F., Halassa, M.M., 2020. Variation of connectivity across exemplar sensory and associative thalamocortical loops in the mouse. eLife 9, e62554. 10.7554/eLife.62554

Müller, E.J., Munn, B., Hearne, L.J., Smith, J.B., Fulcher, B., Arnatkevičiūtė, A., Lurie, D.J., Cocchi, L., Shine, J.M., 2020. Core and matrix thalamic sub-populations relate to spatio-temporal cortical connectivity gradients. NeuroImage 222, 117224. 10.1016/j.neuroimage.2020.117224

Najdenovska, E., Alemán-Gómez, Y., Battistella, G., Descoteaux, M., Hagmann, P., Jacquemont, S., Maeder, P., Thiran, J.-P., Fornari, E., Bach Cuadra, M., 2018. In-vivo probabilistic atlas of human thalamic nuclei based on diffusion- weighted magnetic resonance imaging. Sci Data 5, 180270. 10.1038/sdata.2018.270

Oldham, S., Ball, G., 2023. A phylogenetically-conserved axis of thalamocortical connectivity in the human brain. Nat Commun 14, 6032. 10.1038/s41467-023-41722-8

O’Muircheartaigh, J., Vollmar, C., Traynor, C., Barker, G.J., Kumari, V., Symms, M.R., Thompson, P., Duncan, J.S., Koepp, M.J., Richardson, M.P., 2011. Clustering probabilistic tractograms using independent component analysis applied to the thalamus. NeuroImage 54, 2020–2032. 10.1016/j.neuroimage.2010.09.054

Paquola, C., Vos De Wael, R., Wagstyl, K., Bethlehem, R.A.I., Hong, S.-J., Seidlitz, J., Bullmore, E.T., Evans, A.C., Misic, B., Margulies, D.S., Smallwood, J., Bernhardt, B.C., 2019. Microstructural and functional gradients are increasingly dissociated in transmodal cortices. PLoS Biol 17, e3000284. 10.1371/journal.pbio.3000284

Phillips, J.W., Schulmann, A., Hara, E., Winnubst, J., Liu, C., Valakh, V., Wang, L., Shields, B.C., Korff, W., Chandrashekar, J., Lemire, A.L., Mensh, B., Dudman, J.T., Nelson, S.B., Hantman, A.W., 2019. A repeated molecular architecture across thalamic pathways. Nat Neurosci 22, 1925–1935. 10.1038/s41593-019-0483-3

Roy, D.S., Zhang, Y., Halassa, M.M., Feng, G., 2022. Thalamic subnetworks as units of function. Nat Neurosci 25, 140–153. 10.1038/s41593-021-00996-1

Royer, J., Rodríguez-Cruces, R., Tavakol, S., Larivière, S., Herholz, P., Li, Q., Vos de Wael, R., Paquola, C., Benkarim, O., Park, B., Lowe, A.J., Margulies, D., Smallwood, J., Bernasconi, A., Bernasconi, N., Frauscher, B., Bernhardt, B.C., 2022. An Open MRI Dataset For Multiscale Neuroscience. Sci Data 9, 569. 10.1038/s41597-022-01682-y

Saalmann, Y.B., Pinsk, M.A., Wang, L., Li, X., Kastner, S., 2012. The Pulvinar Regulates Information Transmission Between Cortical Areas Based on Attention Demands. Science 337, 753–756. 10.1126/science.1223082

Salimi-Khorshidi, G., Douaud, G., Beckmann, C.F., Glasser, M.F., Griffanti, L., Smith, S.M., 2014. Automatic denoising of functional MRI data: Combining independent component analysis and hierarchical fusion of classifiers. NeuroImage 90, 449–468. 10.1016/j.neuroimage.2013.11.046

Saranathan, M., Iglehart, C., Monti, M., Tourdias, T., Rutt, B., 2021. In vivo high-resolution structural MRI-based atlas of human thalamic nuclei. Sci Data 8, 275. 10.1038/s41597-021-01062-y

Schaefer, A., Kong, R., Gordon, E.M., Laumann, T.O., Zuo, X.-N., Holmes, A.J., Eickhoff, S.B., Yeo, B.T.T., 2018. Local-Global Parcellation of the Human Cerebral Cortex from Intrinsic Functional Connectivity MRI. Cerebral Cortex 28, 3095–3114. 10.1093/cercor/bhx179

Schiff, N.D., 2008. Central Thalamic Contributions to Arousal Regulation and Neurological Disorders of Consciousness. Annals of the New York Academy of Sciences 1129, 105–118. 10.1196/annals.1417.029

Schmitt, J.E., Lenroot, R., Ordaz, S.E., Wallace, G.L., Lerch, J.P., Evans, A.C., Prom, E.C., Kendler, K.S., Neale, M.C., Giedd, J.N., 2009. Variance Decomposition of MRI-Based Covariance Maps Using Genetically-Informative Samples and Structural Equation Modeling. Neuroimage 47, 56–64. 10.1016/j.neuroimage.2008.06.039

Segall, J.M., Allen, E.A., Jung, R.E., Erhardt, E.B., Arja, S.K., Kiehl, K., Calhoun, V.D., 2012. Correspondence between structure and function in the human brain at rest. Front. Neuroinform. 6. 10.3389/fninf.2012.00010

Sherman, S.M., 2016. Thalamus plays a central role in ongoing cortical functioning. Nat Neurosci 19, 533–541. 10.1038/nn.4269

Sherman, S.M., 2012. Thalamocortical interactions. Current Opinion in Neurobiology, Microcircuits 22, 575–579. 10.1016/j.conb.2012.03.005

Sherman, S.M., Guillery, R.W., 2013. Functional Connections of Cortical Areas: A New View from the Thalamus. MIT Press.

Sherman, S.M., Guillery, R.W., 1998. On the actions that one nerve cell can have on another: Distinguishing “drivers” from “modulators.” Proc. Natl. Acad. Sci. U.S.A. 95, 7121–7126. 10.1073/pnas.95.12.7121

Shine, J.M., Lewis, L.D., Garrett, D.D., Hwang, K., 2023. The impact of the human thalamus on brain-wide information processing. Nat Rev Neurosci 24, 416–430. 10.1038/s41583-023-00701-0

Smallwood, J., Bernhardt, B.C., Leech, R., Bzdok, D., Jefferies, E., Margulies, D.S., 2021. The default mode network in cognition: a topographical perspective. Nat Rev Neurosci 22, 503–513. 10.1038/s41583-021-00474-4

Smith, S.M., Jenkinson, M., Woolrich, M.W., Beckmann, C.F., Behrens, T.E.J., Johansen-Berg, H., Bannister, P.R., De Luca, M., Drobnjak, I., Flitney, D.E., Niazy, R.K., Saunders, J., Vickers, J., Zhang, Y., De Stefano, N., Brady, J.M., Matthews, P.M., 2004. Advances in functional and structural MR image analysis and implementation as FSL. NeuroImage 23, S208–S219. 10.1016/j.neuroimage.2004.07.051

Stough, J.V., Glaister, J., Ye, C., Ying, S.H., Prince, J.L., Carass, A., 2014. Automatic Method for Thalamus Parcellation Using Multi-modal Feature Classification, in: Golland, P., Hata, N., Barillot, C., Hornegger, J., Howe, R. (Eds.), Medical Image Computing and Computer-Assisted Intervention – MICCAI 2014, Lecture Notes in Computer Science. Springer International Publishing, Cham, pp. 169–176. 10.1007/978-3-319-10443-0_22

Su, J.H., Thomas, F.T., Kasoff, W.S., Tourdias, T., Choi, E.Y., Rutt, B.K., Saranathan, M., 2019. Thalamus Optimized Multi Atlas Segmentation (THOMAS): fast, fully automated segmentation of thalamic nuclei from structural MRI. NeuroImage 194, 272–282. 10.1016/j.neuroimage.2019.03.021

Sydnor, V.J., Larsen, B., Seidlitz, J., Adebimpe, A., Alexander-Bloch, A.F., Bassett, D.S., Bertolero, M.A., Cieslak, M., Covitz, S., Fan, Y., Gur, R.E., Gur, R.C., Mackey, A.P., Moore, T.M., Roalf, D.R., Shinohara, R.T., Satterthwaite, T.D., 2023. Intrinsic activity development unfolds along a sensorimotor–association cortical axis in youth. Nat Neurosci 26, 638–649. 10.1038/s41593-023-01282-y

Tourdias, T., Saranathan, M., Levesque, I.R., Su, J., Rutt, B.K., 2014. Visualization of intra-thalamic nuclei with optimized white-matter-nulled MPRAGE at 7T. NeuroImage 84, 534–545. 10.1016/j.neuroimage.2013.08.069

Traynor, C., Heckemann, R.A., Hammers, A., O’Muircheartaigh, J., Crum, W.R., Barker, G.J., Richardson, M.P., 2010. Reproducibility of thalamic segmentation based on probabilistic tractography. NeuroImage 52, 69–85. 10.1016/j.neuroimage.2010.04.024

Tustison, N., Avants, B., 2013. Explicit B-spline regularization in diffeomorphic image registration. Frontiers in Neuroinformatics 7.

Tustison, N.J., Avants, B.B., Cook, P.A., Yuanjie Zheng, Egan, A., Yushkevich, P.A., Gee, J.C., 2010. N4ITK: Improved N3 Bias Correction. IEEE Trans. Med. Imaging 29, 1310–1320. 10.1109/TMI.2010.2046908

Valk, S.L., Xu, T., Margulies, D.S., Masouleh, S.K., Paquola, C., Goulas, A., Kochunov, P., Smallwood, J., Yeo, B.T.T., Bernhardt, B.C., Eickhoff, S.B., 2020. Shaping brain structure: Genetic and phylogenetic axes of macroscale organization of cortical thickness. Sci. Adv. 6, eabb3417. 10.1126/sciadv.abb3417

Valk, S.L., Xu, T., Paquola, C., Park, B., Bethlehem, R.A.I., Vos de Wael, R., Royer, J., Masouleh, S.K., Bayrak, Ş., Kochunov, P., Yeo, B.T.T., Margulies, D., Smallwood, J., Eickhoff, S.B., Bernhardt, B.C., 2022. Genetic and phylogenetic uncoupling of structure and function in human transmodal cortex. Nat Commun 13, 2341. 10.1038/s41467-022-29886-1

Veraart, J., Novikov, D.S., Christiaens, D., Ades-aron, B., Sijbers, J., Fieremans, E., 2016. Denoising of diffusion MRI using random matrix theory. NeuroImage 142, 394–406. 10.1016/j.neuroimage.2016.08.016

Vogel, J.W., Alexander-Bloch, A., Wagstyl, K., Bertolero, M., Markello, R., Pines, A., Sydnor, V.J., Diaz-Papkovich, A., Hansen, J., Evans, A.C., Bernhardt, B., Misic, B., Satterthwaite, T., Seidlitz, J., 2022. Conserved whole-brain spatiomolecular gradients shape adult brain functional organization. 10.1101/2022.09.18.508425

Vos de Wael, R., Benkarim, O., Paquola, C., Lariviere, S., Royer, J., Tavakol, S., Xu, T., Hong, S.-J., Langs, G., Valk, S., Misic, B., Milham, M., Margulies, D., Smallwood, J., Bernhardt, B.C., 2020. BrainSpace: a toolbox for the analysis of macroscale gradients in neuroimaging and connectomics datasets. Commun Biol 3, 103. 10.1038/s42003-020-0794-7

Vos de Wael, R., Royer, J., Tavakol, S., Wang, Y., Paquola, C., Benkarim, O., Eichert, N., Larivière, S., Xu, T., Misic, B., Smallwood, J., Valk, S.L., Bernhardt, B.C., 2021. Structural Connectivity Gradients of the Temporal Lobe Serve as Multiscale Axes of Brain Organization and Cortical Evolution. Cereb Cortex 31, 5151–5164. 10.1093/cercor/bhab149

Waehnert, M.D., Dinse, J., Weiss, M., Streicher, M.N., Waehnert, P., Geyer, S., Turner, R., Bazin, P.- L., 2014. Anatomically motivated modeling of cortical laminae. NeuroImage 93, 210–220. 10.1016/j.neuroimage.2013.03.078

Weiskopf, N., Edwards, L.J., Helms, G., Mohammadi, S., Kirilina, E., 2021. Quantitative magnetic resonance imaging of brain anatomy and in vivo histology. Nat Rev Phys 3, 570–588. 10.1038/s42254-021-00326-1

Wolff, M., Morceau, S., Folkard, R., Martin-Cortecero, J., Groh, A., 2021. A thalamic bridge from sensory perception to cognition. Neuroscience & Biobehavioral Reviews 120, 222–235. 10.1016/j.neubiorev.2020.11.013

Yang, S., Meng, Y., Li, J., Li, B., Fan, Y.-S., Chen, H., Liao, W., 2020. The thalamic functional gradient and its relationship to structural basis and cognitive relevance. NeuroImage 218, 116960. 10.1016/j.neuroimage.2020.116960

Yee, Y., Ellegood, J., French, L., Lerch, J.P., 2024. Organization of thalamocortical structural covariance and a corresponding 3D atlas of the mouse thalamus. NeuroImage 285, 120453. 10.1016/j.neuroimage.2023.120453

Yee, Y., Fernandes, D.J., French, L., Ellegood, J., Cahill, L.S., Vousden, D.A., Spencer Noakes, L., Scholz, J., van Eede, M.C., Nieman, B.J., Sled, J.G., Lerch, J.P., 2018. Structural covariance of brain region volumes is associated with both structural connectivity and transcriptomic similarity. NeuroImage 179, 357–372. 10.1016/j.neuroimage.2018.05.028

Yeo, B.T., Krienen, F.M., Sepulcre, J., Sabuncu, M.R., Lashkari, D., Hollinshead, M., Roffman, J.L., Smoller, J.W., Zöllei, L., Polimeni, J.R., Fischl, B., Liu, H., Buckner, R.L., 2011. The organization of the human cerebral cortex estimated by intrinsic functional connectivity. Journal of Neurophysiology 106, 1125–1165. 10.1152/jn.00338.2011

Zhang, D., Snyder, A.Z., Fox, M.D., Sansbury, M.W., Shimony, J.S., Raichle, M.E., 2008. Intrinsic Functional Relations Between Human Cerebral Cortex and Thalamus. Journal of Neurophysiology 100, 1740–1748. 10.1152/jn.90463.2008

Zheng, W., Zhao, L., Zhao, Z., Liu, T., Hu, B., Wu, D., 2023. Spatiotemporal Developmental Gradient of Thalamic Morphology, Microstructure, and Connectivity fromthe Third Trimester to Early Infancy. J. Neurosci. 43, 559–570. 10.1523/JNEUROSCI.0874-22.2022

